# A cofactor-promiscuous HMGR from the Lyme disease pathogen illuminates diversity in bacterial isoprenoid biosynthesis

**DOI:** 10.64898/2026.07.06.735745

**Authors:** Isaac A. Paddy, Joshua McCausland, Madelyn Frazier, Poulami Chatterjee, Mekedlawit Setegne, Oliv Eidam, Christine Jacobs-Wagner, Laura M. K. Dassama

## Abstract

The Lyme disease pathogen *Borrelia burgdorferi* contains a highly reduced genome lacking many primary metabolic pathways. However, *B. burgdorferi* retains the mevalonate pathway that synthesizes isopentenyl pyrophosphate (IPP), the precursor to the peptidoglycan carrier lipid. While the mevalonate pathway and the enzyme that catalyzes its rate-limiting step (3-hydroxy-3-methyl glutaryl coenzyme A reductase, HMGR) are well studied in vertebrates, little is known about the pathway in *B. burgdorferi* and many pathogenic bacteria. In this work, we reveal that HMGR is a critical metabolic enzyme in *B. burgdorferi*. We demonstrate that loss of HMGR causes morphological defects and muted de novo synthesis of peptidoglycan; these defects are ameliorated by exogenous mevalonate and IPP. Biochemical characterization unveiled the HMGR as a highly unusual cofactor-promiscuous oxidoreductase that functions with both nicotinamide cofactors. Bioinformatics and biochemical characterization uncovered examples of similarly promiscuous HMGRs and revealed a previously unrecognized evolutionary link to cofactor choice. Moreover, structures of the enzyme reveal a highly divergent active site architecture. Together, these findings firmly establish HMGR as an opportunity target for the development of antibacterials for a diderm pathogen while highlighting cofactor promiscuity as an evolutionary acquired feature in HMGRs.

## Introduction

The diderm *Borrelia burgdorferi* is the causative agent of Lyme disease, which is the most common zoonotic illness in certain parts of the world^1^. *B. burgdorferi* has a reduced genome (1,291 protein-encoding genes in the B31 strain compared to > 4,000 in *Escherichia coli*) and lacks essential metabolic pathways such as the TCA cycle and pathways for the de novo synthesis of nucleotides, amino acids, and fatty acids^2–6^. Moreover, the pathogen performs glycolysis with a limited set of carbohydrates^3, 7, 8^ and is incapable of performing fatty acid catabolism. This metabolic minimalism makes the pathogen susceptible to nutrient limitation^9^. Thus, targeting the machineries used to acquire or synthesize essential metabolites may be a powerful strategy to curtail the proliferation of the pathogen.

The mevalonate (MVA) pathway is one of the few conserved metabolic pathways in *B. burgdorferi*^10^. This pathway is found in the three domains of life and converts acetate to isopentenyl pyrophosphate (IPP)^6, 11–14^. IPP is the single isoprene unit that is extended and modified to make isoprenoids such as sterols and the peptidoglycan lipid carrier undecaprenyl-phosphate (C55-P). Beyond these roles, isoprenoids are used to modify proteins via prenylation and are important for a myriad of cellular processes related to cell growth and differentiation^15–17^. Because of these roles in important cellular processes, enzymes catalyzing the formation of IPP are thought to be essential for viability. In eukaryotes, the pathway is best known for its role in cholesterol biosynthesis and is the target of the hypercholesterolemia-treating statin drugs^11^. In bacteria, the MVA pathway has been linked to peptidoglycan synthesis^18, 19^ and post-translational modifications of proteins through hijacking of eukaryotic host machinery^20, 21^. The rate-limiting step of the pathway is the conversion of hydroxymethylglutaryl-CoA (HMG-CoA) to mevalonate, a reaction that is catalyzed by the enzyme HMG-CoA reductase (HMGR). This four-electron reduction requires two molecules of NAD(P)H to reduce the thioester in HMG-CoA to an alcohol.

Attempts to genetically silence key biosynthetic enzymes of the MVA pathway (including HMGR) in other organisms have failed, and the assertion is that the essentiality of the pathway is responsible for its genetic intractability^14^. It is likely that the MVA pathway in *B. burgdorferi* is used for the synthesis of peptidoglycan via C55-P, and there is some evidence hinting at its relevance. *B. burgdorferi* encodes all homologs of the MVA pathway enzymes needed to make IPP in an operon; it also has homologs of genes involved in the synthesis of geranyl pyrophosphate (GPP), farnesyl pyrophosphate (FPP), undecaprenyl pyrophosphate (C55-PP), and undecaprenyl phosphate (C55-P); C55-P is the dedicated lipid carrier for peptidoglycan precursors (Fig. S1)^6, 22^. Published work has shown that cultured *B. burgdorferi* is sensitive to acetate levels, implicating the MVA pathway as the culprit for this sensitivity^14^. Furthermore, a recent in vitro metabolic reconstitution study suggested that the mevalonate pathway is essential for viability in *B. burgdorferi*^10^. We posit that the limited metabolic capability of *B. burgdorferi* heightens its dependence on the existing metabolic pathways. As such, perturbations to the MVA pathway via HMGR might prove lethal to the pathogen.

HMGR enzymes can be grouped into two classes based on protein sequence, with class I enzymes primarily found in eukaryotes and in some archaea and bacteria while class II enzymes are exclusive to bacteria and archaea^23, 24^. Class I HMGRs are mostly tetrameric and exclusively use NADPH as a redox cofactor^24^. On the contrary, class II HMGRs can adopt multiple oligomeric states and can use either NADPH or NADH as cofactors with varying degrees. For example, class II HMGR enzymes from *Burkholderia cenocepacia^25^*and *Pseudomonas mevalonii*^26^ use only NADH while the HMGR from *Enterococcus faecalis*^27^ is functional with NADPH only. While some class II HMGRs have been reported to use both cofactors, they all display a strong preference for one. Amongst these are the HMGR from *Delftia acidovorans*^28^ (prefers NADH), *Archaeoglobus fulgidus*^29^, *Listeria monocytogenes*^30^, *Staphylococcus aureus*^18, 31^, and *Streptococcus pneumoniae*^28^ (NADPH-preferring). Despite these two cofactors being functionally identical, evolution has finely tuned enzyme active sites to select for one; the ability to use both is rare and may have implications for adaptation necessary to survive in different environments during the microbe’s life cycle. Moreover, ambivalent enzymes whose cofactor selectivity can be altered are critical for synthetic biology applications geared at engineering cellular metabolism^32,33^.

In this work we sought to establish the essentiality and role of the HMGR from the metabolically minimal pathogen *B. burgdorferi*. Using a combination of genetic and chemical tools, we asked whether the enzyme is essential for proliferation and interrogated its role in peptidoglycan biogenesis. We further integrated biochemical, structural, and large-scale bioinformatics to dissect the molecular features of the HMGR enzyme from *B. burgdorferi* and assessed how cofactor usage and divergence of active-site architectures vary across bacterial HMGRs.

## Results

### Gene silencing of hmgr leads to severe morphological defects and muted de novo peptidoglycan synthesis in B. burgdorferi

To investigate the relevance of HMGR, we created a CRISPR interference (CRISPRi) shuttle vector^34^, which carries an isopropyl-β-D-1-thiogalactopyranoside (IPTG)-inducible guide RNA targeting bp133 of the *hmgr/bb0685* gene. We introduced this resulting vector into *B. burgdorferi* B31 strain K2 to create strain CJW_Bb662. In the absence of IPTG, the culture grew to a density comparable to that of the wild-type strain (∼ 10^8^ cells/mL 5 days post-inoculation, **Fig. S2B**). Upon addition of ITPG, phase-contrast microscopy revealed drastic membrane blebbing and reduced mobility for cells incubated with IPTG unlike those grown in the absence of IPTG (**Fig. 1A**, **Movies S1 and S2**). To examine whether the observed morphological defects arise from the loss of *hmgr* expression and not polar effects on neighboring genes, we performed quantitative real-time PCR to amplify transcripts of *hmgr* and downstream genes: *bb0686*, *bb0687*, and *bb0688.* The results confirmed that CRISPR interference was largely specific to *hmgr* (p < 0.05 using one-way ANOVA with post-hoc Tukey test) (**Fig. S2C**).

**Fig. 1.**
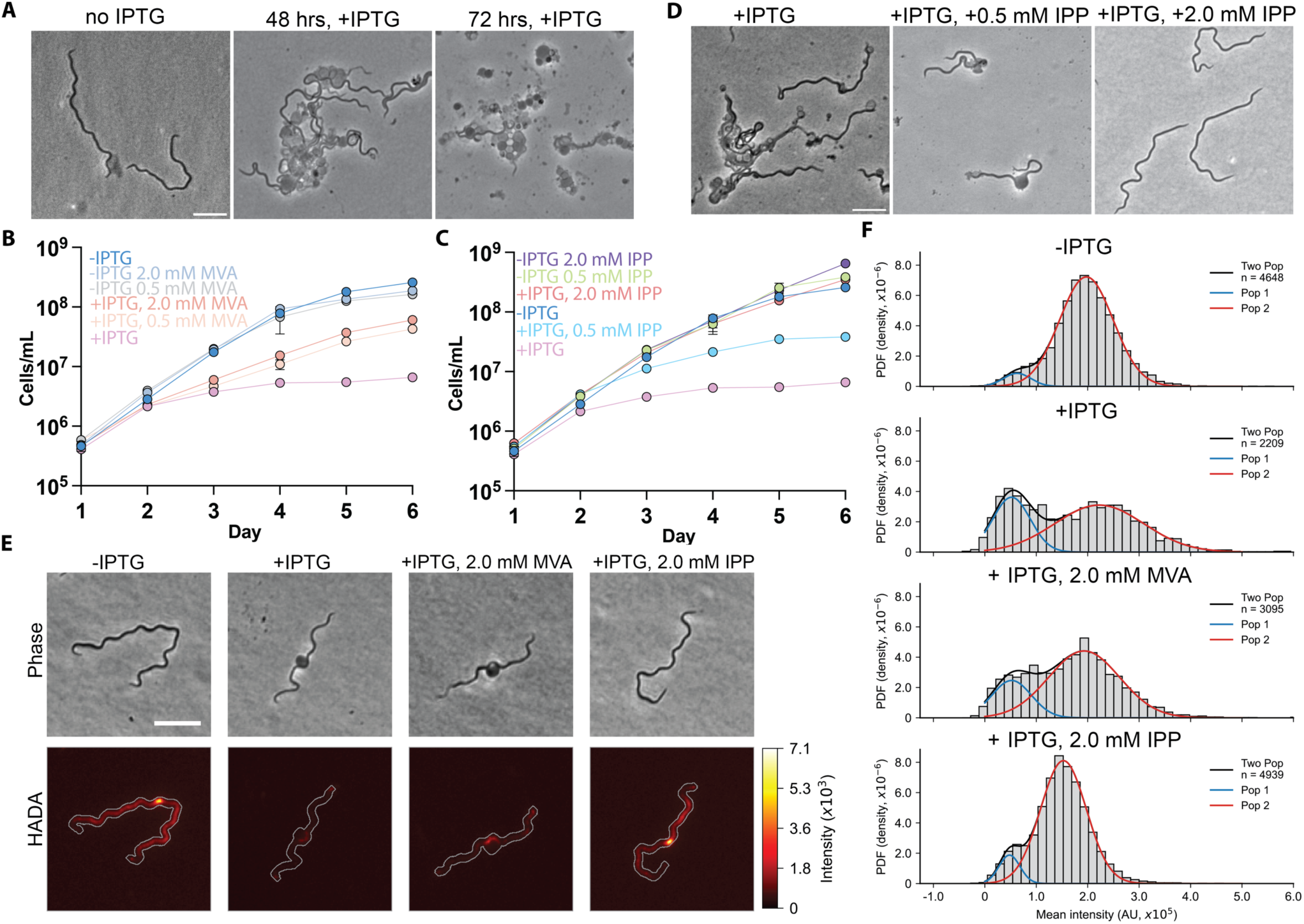
Phenotypes associated with *hgmr* silencing in *B. burgdorferi* by CRISPR interference. **A**) Representative images of cells taken from cultures with or without IPTG. Scale bar: 5 µm. **B**) Growth curve of *B. burgdorferi* (strain CJW_Bb662) growing in the absence (uninduced) or presence (induced) of 100 µM IPTG to silence *hmgr*. For some conditions, the cultures were supplemented with 0, 0.5, and 2.0 mM MVA. **C**) Same as panel B but with or without supplementation of 0, 0.5, and 2.0 mM IPP. **D**) Representative images of CJW_Bb662 cells from cultures grown in the presence of the indicated compounds. Scale bar: 5 µm. **E**) Representative images of cells incubated with HADA. Each image in the HADA channel was normalized to the same indicated minimum and maximum values of HADA signal intensity in arbitrary units (au). Each fluorescent component of the merged images is auto-contrasted. The scale bar: 5 µm. **F)** Distributions of single-cell HADA intensities from all four indicated conditions. The black lines show the Gaussian fit on the two populations (pop 1 and pop 2), whereas the blue and red lines show the contribution of each individual population. The n value is indicated. They represent three pooled biological replicates (shown separately in Fig. S2D).

MVA is the direct product of the enzyme’s reaction with HMG-CoA. To determine whether MVA in the growth medium ameliorates the effect of *hmgr* silencing, we measured the growth of cultures without IPTG, with IPTG added to induce HMGR depletion, and with IPTG and MVA. These data revealed a partial reversal of the growth defect with the addition of 500 μM MVA but even at 2 mM MVA, the HMGR-depleted cells still have a substantial growth defect (**Fig. 1B**). The same experiments were done but with IPP supplementation, which partially restored cell growth at 500 μM IPP and fully reversed the phenotype at 2 mM IPP (**Fig. 1C**). Representative phase-contrast microscopy revealed that IPP supplementation partially (0.5 mM) or fully (2 mM) rescued the morphological defects of the HGMR-depleted cells (**Fig. 1D)**. The fact that IPP but not MVA is able to fully rescue the phenotypes associated with HMGR depletion may suggest a difference in uptake between the two metabolites, although we cannot exclude the possibility of a mild silencing effect on the genes downstream of *hmgr* contributes to the observed defects.

Given that IPP is an essential precursor to C55-P, we hypothesized that HMGR depletion impacts peptidoglycan synthesis. To detect changes in *de novo* PG biosynthesis, we used the fluorescent D-alanine analog 7-hydroxycoumarin 3-carboxylic acid linked to 3-amino-D-alanine (HADA)^35^, which mimics the terminal D-alanine residue of the peptidoglycan backbone. When added exogenously, *de novo* peptidoglycan synthesis results in the incorporation of HADA to the *B. burgdorferi* peptidoglycan^36^. Fluorescent imaging revealed reduced HADA labeling in IPTG-induced cells that display the blebbing phenotype characteristic of the HMGR depletion (**Fig. 1E**). Single-cell analysis revealed that HMGR depletion results in two populations of cells, and that one population has substantially reduced fluorescence (**Figs. 1F**, **S2D, Table S1**). The addition of IPP largely rescued this phenotype, as did MVA to a lesser degree (**Figs. 1E, F, S2D**). These data are consistent with a growth reduction associated with a defect in PG biosynthesis and provide the first direct observation of muted peptidoglycan synthesis upon loss of *hmgr*.

### B. burgdorferi HMGR is promiscuous with redox cofactors

To interrogate the activity of *B. burgdorferi* HMGR, we produced the protein recombinantly in *E. coli* as a fusion of maltose-binding protein to aid its solubility (**Figs. S3, S4A**). Following affinity chromatography and removal of the solubility tag, pure HMGR was obtained as a dimer; addition of cofactor and substrates did not alter its oligomerization (**Fig. 2A and Figs. S4B-F**). Previous studies with an MBP-fusion of *B. burgdorferi* HMGR revealed that the enzyme can mediate the oxidation of NADPH^14^. Using the established absorbance-based assay that measures cofactor oxidation via a reduction in its 340 nm peak, we determined that the enzyme indeed functions with NADPH, exhibiting a *K*_M_ of 7.66 ± 1.34 μM for HMG-CoA (**Figs. 2B, C**).

**Fig. 2.**
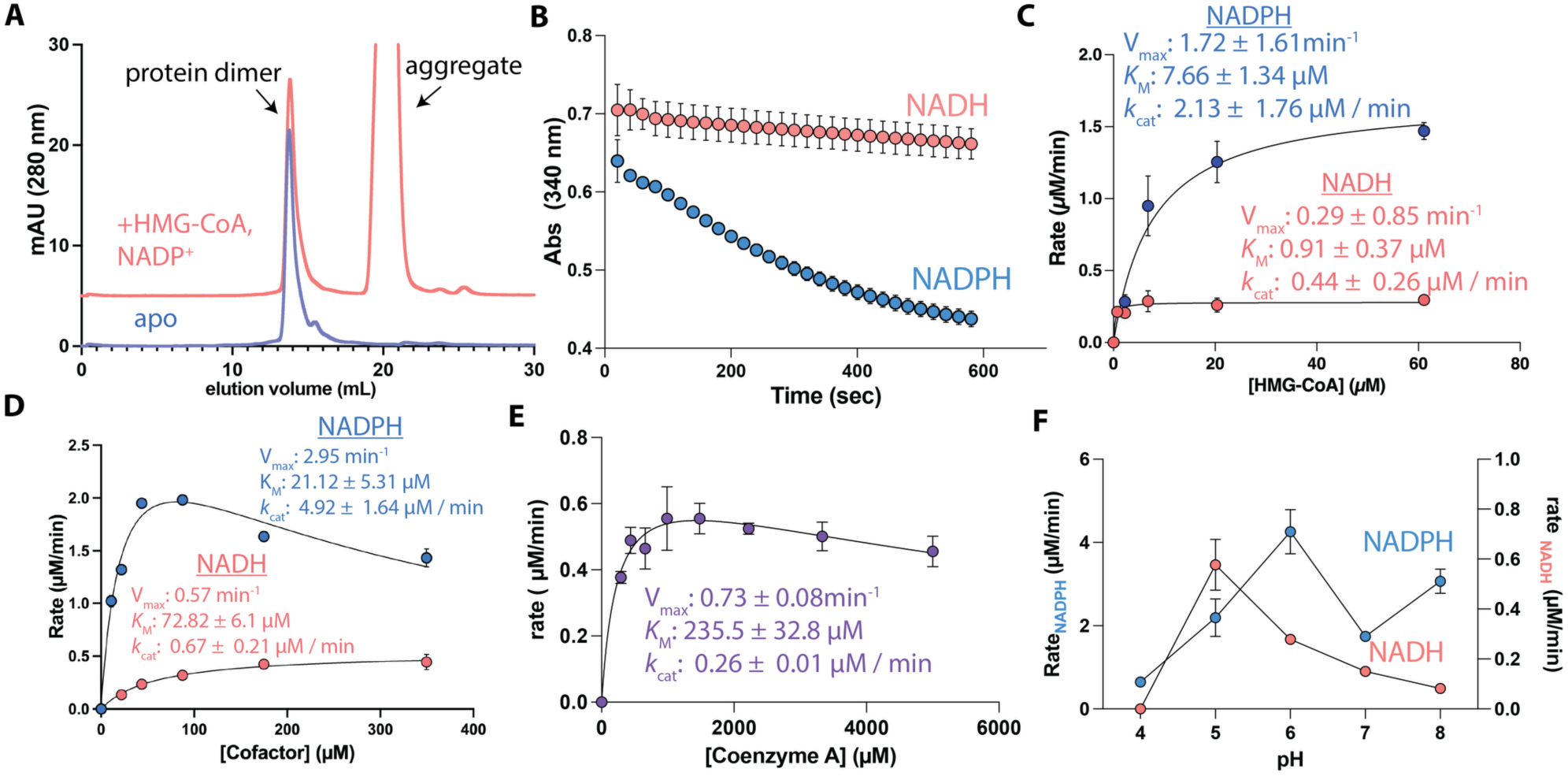
**Biochemical characterization of HMGR from *B. burgdorferi***. **A**) Size-exclusion chromatography of HMGR (blue) and with the addition of 100 µM HMG-CoA and NADP^+^(pink). **B)** Representative time vs absorbance plot of enzymatic reaction using 125 µM of HMG-CoA, 600 µM of cofactor, and 500 nM enzyme. Steady-state kinetics of HMGR as a function of substrate (**C**) and cofactor (**D**). Concentrations of HMG-CoA varied from 0-500 µM while that of the cofactor ranged 0 – 500 µM NAD(P)H. All assays used 500 nM HMGR, 5 mM β-ME, 150 mM NaCl, 10 % glycerol in 20 mM HEPES 7.5 and were performed at 30 ^∘^C. **E)** Steady-state kinetics monitoring the enzyme’s oxidation of NAD^+^ in the presence of mevalonate and CoA. Assays were performed with 195-5000 µM CoA, 5 mM mevalonate and NADP^+^, and 2.4 µM enzyme at 30 ^∘^C in 20 mM HEPES 7.5 and 150 mM NaCl. **F**) Activity *vs* pH profile of HMGR activity in assays with NADPH or NADH.

While most oxidoreductases are functional with a single cofactor, a handful of bacterial enzymes are known to display a low level of activity with both cofactors^26, 30^. To determine the cofactor specificity of the *B. burgdorferi* enzyme, we performed the same assays with NADH. Whereas the enzyme functioning with NADPH displays a *k_cat_* of 4.92 ± 1.64 min^-1^, this value is lower but detectable with NADH (0.67 ± 0.21 min^-1^). The respective *K*_M_ values are also different, with that of NADPH being ∼ 3-fold lower than that of NADH (21.12 ± 5.31 vs 72.82 ± 6.1 μM, **Figs. 2C, D,** and **Table 1**). Unlike some HMGRs, the enzyme also prefers to catalyze the conversion of HMG-CoA to MVA rather than the reverse, given the much lower *k_cat_* and *K*_M_ values for that reaction (**Fig. 2E**).

**Table 1.**
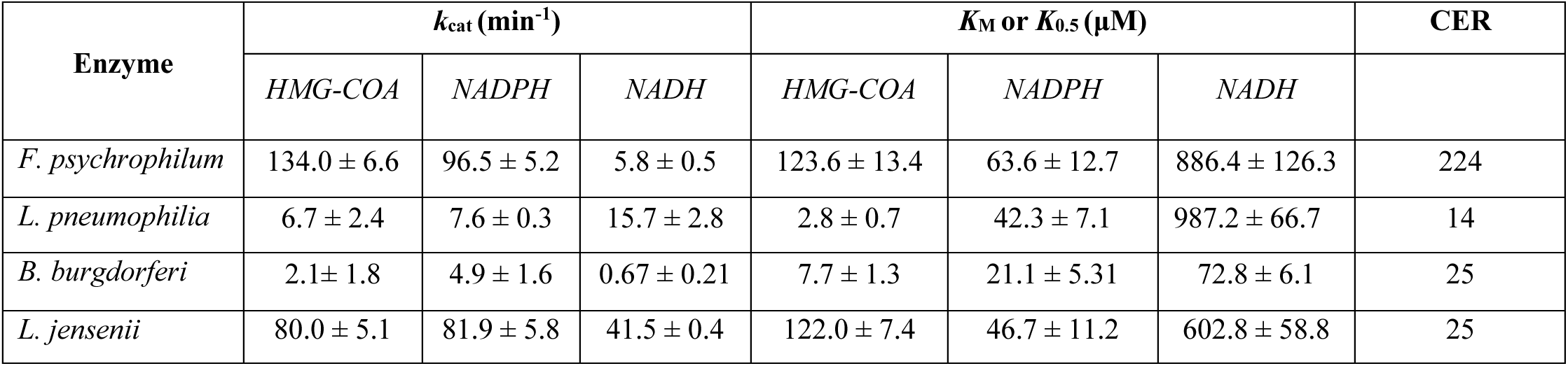
Steady-state kinetics of select HMGR homologs in the “undefined” branches.

|  |  |  |  |  |  |  |  |
| --- | --- | --- | --- | --- | --- | --- | --- |
| Enzyme | $k_{\text{cat}}$ (min<sup>-1</sup>) | | | $K_M$ or $K_{0.5}$ (μM) | | | CER |
|  | *HMG-COA* | *NADPH* | *NADH* | *HMG-COA* | *NADPH* | *NADH* |  |
| *F. psychrophilum* | 134.0 ± 6.6 | 96.5 ± 5.2 | 5.8 ± 0.5 | 123.6 ± 13.4 | 63.6 ± 12.7 | 886.4 ± 126.3 | 224 |
| *L. pneumophila* | 6.7 ± 2.4 | 7.6 ± 0.3 | 15.7 ± 2.8 | 2.8 ± 0.7 | 42.3 ± 7.1 | 987.2 ± 66.7 | 14 |
| *B. burgdorferi* | 2.1 ± 1.8 | 4.9 ± 1.6 | 0.67 ± 0.21 | 7.7 ± 1.3 | 21.1 ± 5.31 | 72.8 ± 6.1 | 25 |
| *L. jensenii* | 80.0 ± 5.1 | 81.9 ± 5.8 | 41.5 ± 0.4 | 122.0 ± 7.4 | 46.7 ± 11.2 | 602.8 ± 58.8 | 25 |

Because the measured *k_cat_* values for the two cofactors differ by 7-fold but the *K*_M_ values are only 3-fold different, we used the catalytic efficiency (*k*_cat_/*K*_M_) ratio (CER) for NADPH/NADH to assess the enzyme’s cofactor specificity^37^. This value for the *B. burgdorferi* HMGR is 25, which is much lower than that reported for the majority of bacterial HMGRs that have CERs of ∼280, implying that they are highly selective for one cofactor over the other^26, 28, 30^. To be certain that the activity detected with NADH was not due to NADPH contamination of the stock, we used the same NADH solutions in reactions with the human HMGR. Consistent with published reports, that enzyme displayed no activity with NADH but was functional with NADPH (**Fig. S5**). Together, these findings suggest that the *B. burgdorferi* HMGR prefers NADPH but can also function with NADH to a degree that is rare amongst HMGRs.

Because the proposed reaction mechanism of HMGRs require proton transfer from an active site glutamic acid (**Fig. S6**), we asked whether the reaction rates with NADH or NADPH would be impacted by the pH. We therefore tested the enzyme activity at pH values that ranged between 4 and 9. These data revealed the enzyme displays peak activity at pH 5 when working with NADH and pH 6 when the reaction uses NADPH (**Fig. 2F**), suggesting that the choice of cofactor influences the catalysis and might signal that slightly different reaction mechanisms are employed when using NADH vs. NADPH.

### Evolutionary link to choice of redox cofactor in HMGRs

While a few bacterial HMGRs have previously been studied, there is poor understanding of the sequence diversity across the family and how that might relate to functional differences. Our investigations with the *B. burgdorferi* HMGR revealed a remarkable ability to use both NADH and NADPH. To investigate the sequence differences amongst bacterial HMGRs that could explain the observed cofactor promiscuity, we constructed a protein sequence similarity network (SSN) based on a BLAST search using the *B. burgdorferi* sequence (**Fig. 3A**). The SSN was filtered for bacterial sequences, resulting in 6,924 nodes and 1,043,350 edges; each node represents a unique protein sequence or sequences that share 100% sequence identity. This network is the first to show all annotated HMGR homologs in bacteria; highlighted are biochemically characterized HMGRs such as those from *Pseudomonas mevalonii*, *Burkholderia cenocepacia*, *E. faecalis*, and more (**Fig 3A**). Among these sequences, *B. burgdorferi* HMGR is separated from the rest of the network in a cluster containing 41 sequences of other *Borrelia* species and uncharacterized spirochetes. This suggests that there are sequence differences from other bacterial homologs that may translate to structural and functional differences.

**Fig. 3.**
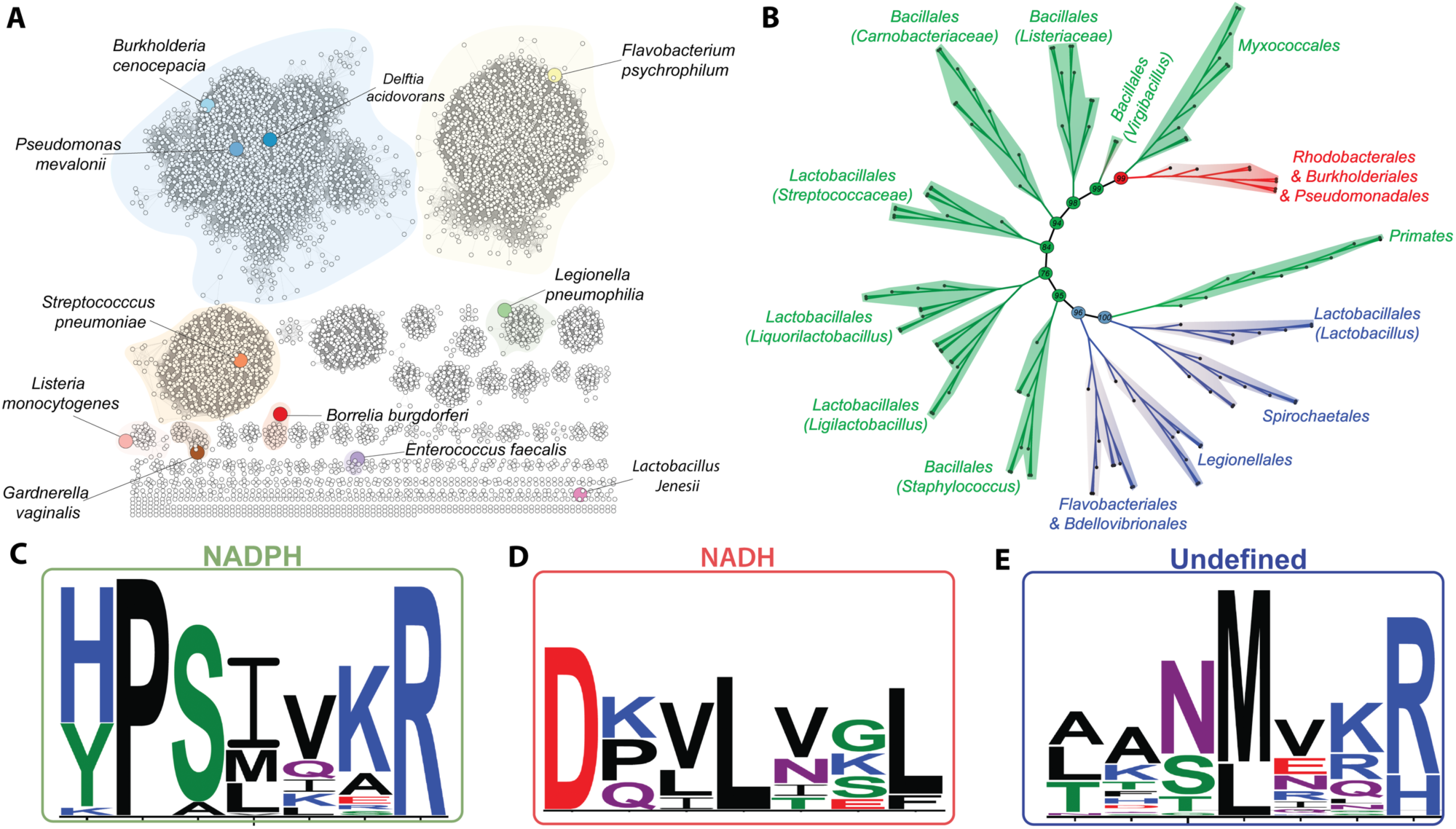
Protein sequence divergence in bacterial HMGRs. **A**) Protein sequence similarity network (SSN) of bacterial HMGRs depicting its size and functional clustering. The SSN was generated via the Enzyme Function Initiatives-Enzyme Similarity tool (EFI-EST) and visualized in Cytoscape. The SSN was generated by employing a single sequence blast of *B. burgdorferi* HMGR and tailored so that the nodes represent sequences with 100% identity, an e-value of 1×10^-5^, and an alignment score of 125. There are 6,924 nodes. Colored in are representative organisms in which a HMGR homolog has been characterized. **B)** Rooted maximum likelihood phylogenetic tree. The tree contains 210 sequences from the 14 largest clusters from the SSN, using eukaryotic HMGRs as an outgroup. The tree was computed with the IQ-tree software and visualized using Figtree. **C-E)** Sequence LOGO of the cofactor helix taken from all sequences used to generate the phylogenetic tree.

To further interrogate these differences, we generated a phylogenetic tree that tracks potential evolutionary relationships between the *B. burgdorferi* HMGR and other bacterial homologs. The maximum likelihood rooted tree contains 140 sequences from the 14 largest clusters in the SSN and uses HMGR sequences from vertebrates as an outgroup (**Fig 3B**). The tree segregates into ten major branches. The first two branches contain only uncharacterized bacterial homologs including the sequences from *B. burgdorferi*, *Legionellae*, *Flavobacteria*, and *Lactobacilli*. Following that are branches with HMGRs from all gram-positive organisms, including a structurally characterized homolog from *S. pneumoniae*^28^; the last branch clade contains homologs from gram-negative bacteria that include structurally characterized enzymes from *P. mevalonii*^38^, *B. cenocepacia*^39^, and *D. acidovorans*^40^.

One notable observation is that the sequences on the phylogenetic tree clustered according to their reported cofactor usage. Miller and Kung^28^ characterized an amino acid motif specific to bacterial HMGRs that are important for binding and positioning (called the “cofactor” helix, which precedes the more established cofactor binding domain that is present in all HMGRs). On the cofactor helix, Dx(V/L)xxxL is found in NADH-binding enzymes while (H/Y)xSxxxR is observed in NADPH-binding HMGRs^28^. Indeed, experimental characterization of many bacterial HMGRs supports Miller’s proposition, and enzymes with these verified or predicted cofactor preferences show a clustering that suggests an evolutionary link to cofactor usage. Surprisingly, HMGR sequences occupying the first 2 branches of the phylogenetic tree deviate from both motifs. These include the aforementioned HMGRs from *B. burgdorferi*, *Legionellae*, *Flavobacteria*, and *Lactobacilli*; we have thus termed these branches the “undefined” clade. To investigate the degree of sequence conservation at the cofactor helix of all HMGRs, we generated a sequence logo of the helix. This multisequence alignment revealed a remarkable sequence conservation, with substantial conservation at position 4 of the motif. Whereas the NADH preferring enzymes all encode a Leu at that position and the NADPH enzymes contain Leu/Ile/Met, the enzymes in the undefined branches encode either Met/Leu (**Fig. 3C-E**). We reasoned that this conservation at position 4 was missed in previous studies that analyzed far fewer sequences. It was also obvious from the logo of the undefined branches that these proteins had amino acids important for functioning with NADH (e.g., the conservation of Met/Leu in position 4) and NADPH (presence of Arg at position 7). We hypothesized that these proteins might function with both cofactors.

### Cofactor selectivity of bacterial HMGR homologs in the “undefined” clade

To probe cofactor promiscuity within the “undefined” clade of the phylogenetic tree, we recombinantly produced a HMGR homolog from each major branch. The proteins from *Legionella pneumophilia*, *Flavobacteria psychrophilum*, and *Lactobacillus jensenii* were purified to homogeneity, with size exclusion chromatography revealing that all but the *L. pneumophilia* homolog are dimeric; this homolog exists in multiple oligomeric states with the minimum being a dimer (**Fig. 4A-C**). Next, we used the dimer versions of each enzyme and the absorbance-based assay previously described to determine the steady-state kinetic parameters of their reaction with varying concentrations of HMG-CoA (**Table 1**, **Fig. 4D-F**). Unlike the *B. burgdorferi* HMGR, these enzymes all of the homologs exhibited a degree of cooperativity (see Hill coefficients in **Fig. 4D-F**). Furthermore, while all preferred NADPH, they still exhibited low level of activity with NADH. To determine whether this amounted to the level of promiscuity detected with the *B. burgdorferi* enzyme, we calculated their CERs when operating with NADPH *vs*. NADH. This revealed that *L. pneumophilia* and *L. jensenii* HMGRs have CERs of 14 and 25, respectively; these values are similar to the CER of 25 determined for the *B. burgdorferi* enzyme. Conversely, the CER of *F. psychrophilum* HMGR is 224 and on par with values calculated for enzymes that are more specific for one cofactor. As such, enzymes from 3 of the 4 nested branches of the undefined clade display a degree of cofactor promiscuity that is rare amongst HMGRs and more broadly, oxidoreductases (**Table 1**).

**Fig. 4.**
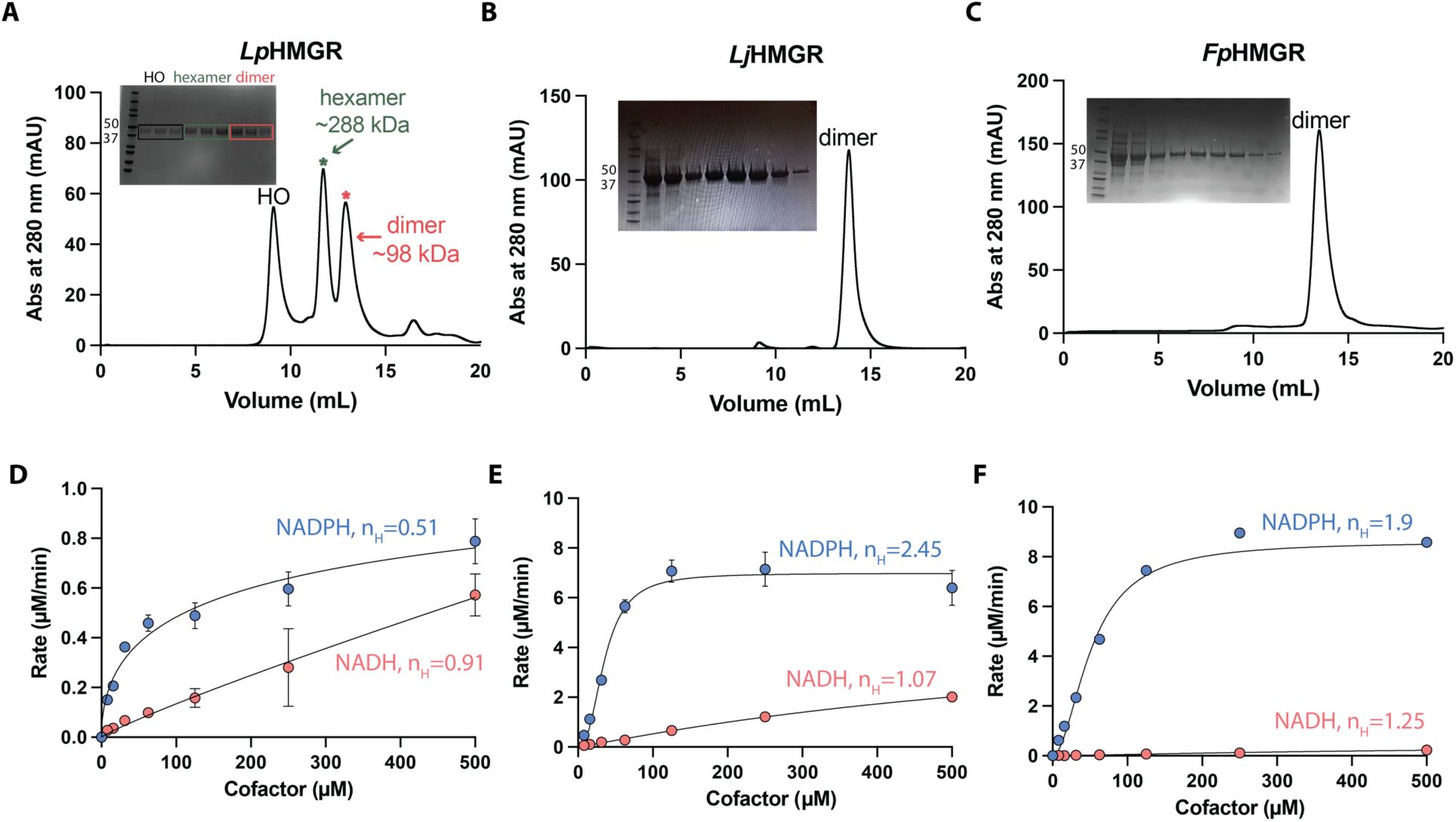
Biochemical characterization of “undefined” HMGRs. A-C) Size exclusion chromatograms (Superdex 200 Increase 10/300 GL) with accompanying SDS-PAGE gels of HMGR homologs from *L. pneumophilia* (*Lp*HMGR), *L. jensenii* (*Lj*HMGR), and *F. psychrophilium* (*Fp*HMGR) respectively. **D-F**) Steady-state kinetics of the HMGRs. Assay conditions were 0.5 µM enzyme, 200 µM HMG-CoA, 0 – 1mM NAD(P)H, in 20 mM HEPES buffer pH 7.5, 5 mM β-ME, 150 mM NaCl, 10 % glycerol. All curves were fit with the allosteric sigmoidal equation provided in the Methods section and have Hill coefficients (n_H_) reported on the figures.

### Statins are poor inhibitors of the B. burgdorferi HMGR

Statins are competitive inhibitors of the human HMGR^41^, disabling the enzyme with high potency and displaying low nanomolar IC_50_ values (**Table S2**). However, statins (**Fig. S7A**), which bind to the HMG-CoA pocket of HMGRs, have been shown to be weak inhibitors of the bacterial homologs^42, 43^. While it has been reported that statins reduce the growth of *B. burgdorferi* and lower the in vitro activity of the MBP-HMGR fusion^14, 44^, kinetic parameters of this inhibition remain unknown. Enzyme activity assays in the presence of free-acid forms of lovastatin, simvastatin, fluvastatin and atorvastatin showed that all statins have IC_50_ values greater than 100 μM, indicating that they are weak inhibitors of *B. burgdorferi* HMGR (**Table S2**, **Fig. S7B**). Similarly, phenyl sulfonamides reported to inhibit a bacterial HMGR^45, 46^ did not alter the activity of the *B. burgdorferi* enzyme (**Fig. S7A, B**). Finally, the addition of statins to the *B. burgdorferi* in vitro cultures did not significantly alter the growth of the pathogen (**Fig. S7C**), unlike the antibiotic streptomycin used here as a control. As such, it is unlikely that statins inhibit HMGR to an appreciable level. The absence of inhibition by the statins hint that, despite the resemblance of HMGRs across species, subtle active site differences dictate the effectiveness of inhibitors.

### Structures of B. burgdorferi HMGR provide insights into its engagement with ligands

To better understand the active site architecture of *B. burgdorferi* HMGR and to additionally gain insights into how potent and selective inhibitors might be identified, we obtained crystals of purified enzyme that diffracted to a resolution of 2.71 Å (**Fig. 5A**, **Table 2**). The *B. burgdorferi* enzyme crystallized as a homodimer, with clear electron density that allowed accurate modeling of most residues in the dimer. Each monomer of HMGR contains a large substrate binding domain for HMG-CoA (**Fig. S8A, B**), small cofactor binding domain that binds NAD(P)H through a non-traditional Rossman fold (hereafter referred to as the cofactor domain) and cofactor helix (a helix that precedes the domain and is unique to bacterial HMGRs, **Fig. S8C**), a three-helical bundle C-terminal domain (residues 372 - 431) that showed discontinuous electron density and for which some residues were not modeled in either chain (**Fig. S8D**), and an intertwined N-terminal domain containing the dimerization motif with residues Glu 54 - Ile 63 (**Fig. S8E**). Despite the protein being crystallized without exogenous ligands, electron density at the active site of each monomer could be modeled as CoA (**Fig. 5A**, **Fig. S9A**). The structure revealed that Lys 97 and Arg 17 form direct hydrogen bonds (or electrostatic interactions) with the negatively charged 2’-phosphate, with distances of 2.8 and 3.0 Å, respectively; whereas Ser 87 forms a hydrogen bond with the amide group of the CoA ligand with a distance of 3.3 Å (**Fig. 5A**).

**Fig. 5.**
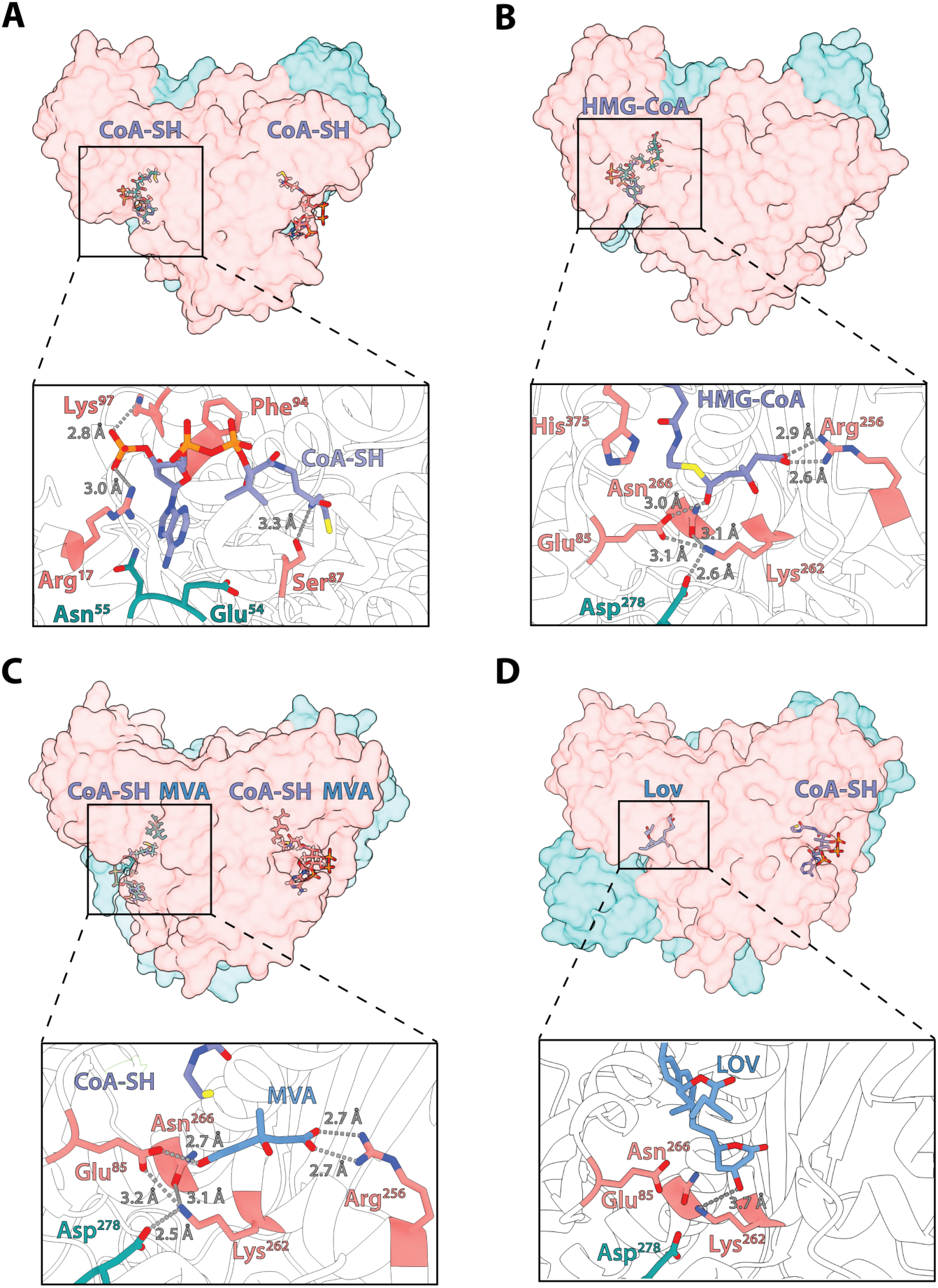
Crystallographic structures of B. burgdorferi HMGR. **A)** Structure of *B. burgdorferi* HMGR with CoA-SH (purple) showing stabilizing interactions in the active site with Arg 17, Ser 87, Lys 97, Phe 94 colored in pink of one monomer and Asn 55 and Glu 54 residues in teal of the second monomer. **B)** Structure of *B. burgdorferi* HMGR with HMG-CoA (purple) showing interactions of the HMG moiety at the active site. Glu 85, Arg 256, Lys 262, Asn 266, and His 375 are shown in pink of one monomer, while Asp 278 from the second monomer is depicted in teal. Red dashes represent potential hydrogen bonds and their respective distances in Å. **C)** Structure of *B. burgdorferi* HMGR with mevalonate (MEV, blue) and CoA-SH (purple). Interacting residues Glu85, Arg 256, Lys 262, and Asn 266 in one monomer are shown in pink while Asp 278 from the second monomer is depicted in teal. **D)** Structure of *B. burgdorferi* HMGR with lovastatin (blue) and CoA-SH (purple), again showing Glu 85, Lys 262, Asn 266 in one monomer in pink while Asp 278 from the second monomer in teal.

**Table 2.** Data collection and refinement statistics (molecular replacement) for *B. burgdorferi* HMGR.

|  | CoA-bound-<br>*Bb*HMGR<br>(PDB: 9ZW8) | HMG-CoA-bound –<br>*Bb*HMGR<br>(PDB: 9ZW6) | Mevalonate-CoA-bound-*Bb*HMGR<br>(PDB: 9ZW9) | Lovastatin-CoA-bound-<br>*Bb*HMGR<br>(PDB: 9ZW7) |
| --- | --- | --- | --- | --- |
| **Data collection** |  |  |  |  |
| Space group | P 21 21 21 | P 21 21 21 | P 21 21 21 | P 21 21 21 |
| Cell dimensions |  |  |  |  |
| *a*, *b*, *c* (Å) | 54.86, 108.87, 139.69 | 54.54, 108.80, 139.62 | 54.44, 109.17, 139.90 | 109.46, 139.92, 54.58 |
| $\alpha$, $\beta$, $\gamma$ (°) | 90.00, 90.00, 90.00 | 90.00, 90.00, 90.00 | 90.00, 90.00, 90.00 | 90.00, 90.00, 90.00 |
| Resolution (Å) | 29.73 - 2.71<br>(2.75 - 2.71) | 24.86 - 2.55<br>(2.61 - 2.55) | 24.89 - 2.20<br>(2.23 - 2.20) | 28.74 - 2.30<br>(2.33 - 2.30) |
| *R*<sub>merge</sub> | 0.1274 (0.9731) | 0.1831 (1.469) | 0.06165 (0.815) | 0.0813 (1.038) |
| *I* / σ*I* | 7.50 (1.13) | 7.51 (1.19) | 13.03 (1.47) | 9.30 (1.50) |
| Completeness (%) | 99.59 (99.89) | 98.66 (97.68) | 98.99 (99.36) | 98.46 (99.73) |
| CC<sub>1/2</sub> (%) | 0.994 (0.577) | 0.825 (0.337) | 0.999 (0.621) | 0.997 (0.612) |
| **Refinement** |  |  |  |  |
| Resolution (Å) | 29.73 - 2.71 | 24.86 - 2.55 | 24.89 - 2.20 | 28.74 - 2.30 |
| No. reflections |  |  |  |  |
| Total | 178734 (6164) | 152497 (8261) | 367069 (14213) | 283413 (11016) |
| Unique | 49262 (1851) | 50333 (3147) | 92897 (3730) | 81141 (3227) |
| Free | 1777 (7.58%) | 2000 (7.14%) | 1747 (4.04%) | 1750 (4.59%) |
| *R*<sub>work</sub> | 0.1993 (0.2346) | 0.2107 (0.2398) | 0.2029 (0.2366) | 0.2086 (0.2466) |
| *R*<sub>free</sub> | 0.2589 (0.3232) | 0.2643 (0.2934) | 0.2503 (0.2957) | 0.2627 (0.3108) |
| No. atoms |  |  |  |  |
| Protein | 6440 (2 copies) | 6417 (2 copies) | 6432 (2 copies) | 6417 (2 copies) |
| Ligand/ion | 96 | 116 | 116 | 108 |
| Water | 45 | 63 | 253 | 119 |
| *B*-factors |  |  |  |  |
| Protein | 59.77 | 49.56 | 49.01 | 53.66 |
| Ligand/ion | 63.94 | 58.57 | 51.28 | 56.71 |
| Water | 60.07 | 49.61 | 49.07 | 50.10 |
| R.m.s. deviations |  |  |  |  |
| Bond lengths (Å) | 0.009 | 0.004 | 0.008 | 0.004 |
| Bond angles (°) | 1.14 | 0.65 | 0.90 | 0.67 |

\*Values in parentheses are for the highest-resolution shell.

The active site of HMGRs is buried at the dimer interface, with HMG-CoA and NAD(P)H binding in a “V”-like shape. The reactive groups of both the substrate and cofactor point towards the middle of the “V”, leaving the remainder of the molecules to extend away from the active site. To determine whether the *B. burgdorferi* enzyme has a similar active site arrangement, we co-crystallized the “as purified” enzyme with HMG-CoA (**Fig. 5B**, **Table 2**) and MVA (**Fig. 5C**, **Table 2**). These yielded crystals that diffracted to 2.55 Å and 2.20 Å, respectively and showed HMG-CoA and MVA occupying only one of the two active sites. The substrate-bound structure revealed that many of the interactions that the protein forms with CoA are also present with the CoA portion of HMG-CoA (**Fig. S9B**). Additionally, the HMG-CoA moiety forms a direct hydrogen bond interaction with Glu 85 with a distance of 3.0 Å (**Fig. 5B**); Glu 85 also is engaged in a hydrogen bonding network with Lys 262, Asn 266, and Asp 278 of the opposite monomer (**Fig. 5B**). Finally, Arg 256 forms a salt bridge with the carboxyl group at the end of the HMG-CoA moiety (**Fig. 5B**). This structure revealed that the substrate forms extensive contacts with the protein.

The structure with MVA bound to the enzyme surprisingly showed density of CoA in both monomers (**Fig. 5C, S9C**), suggesting that both products of the reaction occupy the active site at the same time. In these structures, CoA occupies the same location as HMG-CoA with key interactions formed between the adenosine ring and residues Arg 17 and Lys 97. The MVA retains the key hydrogen bonding network formed with Glu 85, Lys 262, Asn 266 Asp 278, and the salt bridge between the carboxyl and Arg 256.

Despite the weak inhibition by statins, we solved the structure of *B. burgdorferi* HMGR bound to the pro-drug form of lovastatin (2.30 Å, **Table 2**). In this structure, one of the active sites contained density for lovastatin while the other contained density that is best fit with CoA. The cyclized HMG moiety of the lactone form of lovastatin makes a single polar interaction with Lys 262 at the active site of *B. burgdorferi* HMGR (**Fig. 5D**, **Fig. S9D**). In contrast, simvastatin bound to the human HMGR forms a minimum of 7 polar contacts^47^ (**Fig. 6A**). While the lovastatin bound to *B. burgdorferi* HMGR is in the inactive lactone form, it is noteworthy that amino acids at the active site that would be available to engage with the free-acid form are mostly absent (**Fig. 6B**). This suggests that statins can bind to the active site of the *B. burgdorferi* enzyme but the absence of polar interactions with the protein prevents them from acting as potent inhibitors. Importantly, given that hydrogen bonds can contribute ∼ 2 kcal/mol to the Gibbs free energy change of binding^48^, a difference of 6 hydrogen bonds can equate to more than 400× increase in K_D_ for the inhibitor. This implies that effective inhibitors for the *B. burgdorferi* HMGR will require structures drastically distinct from statins that also better leverage the highly hydrophobic active site of the enzyme.

**Fig. 6.**
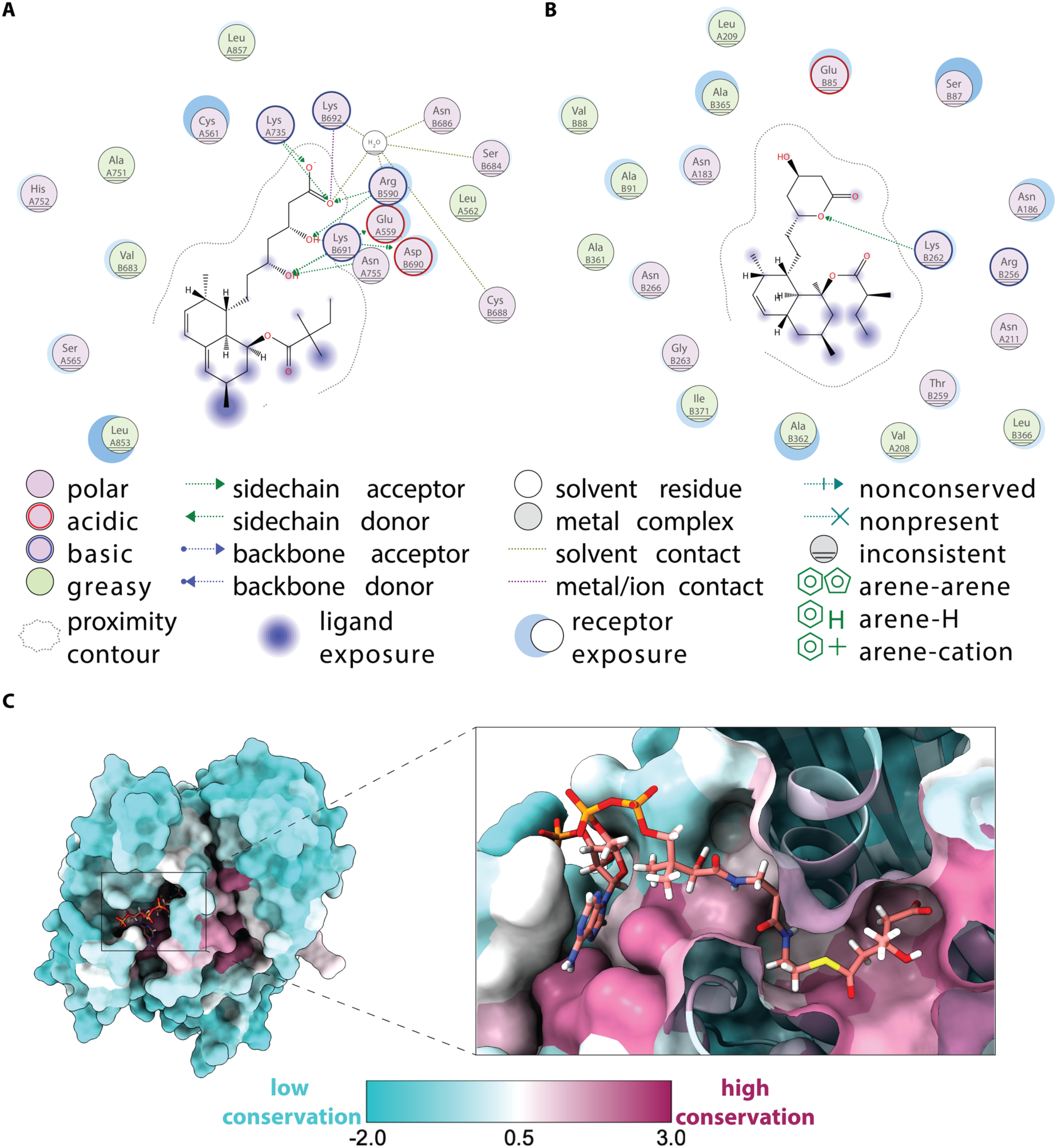
Active site architecture and conservation in HMGRs. a, **b)** Two-dimensional interaction maps of the *H. sapiens* HMGR bound to simvastatin (**A**) the *B. burgdorferi* HMGR bound to lovastatin (**B**). Interaction maps were made and visualized in MOE 2024.06. **c)** HMGR bound to HMG-CoA at the active site with the conservation of active site residues mapped using the sequences generated from the phylogenetic tree. A score of 3 (pink) indicates high sequence conservation at that position, whereas a score of −2.0 (blue) indicates low sequence conservation at that position. Visualized in ChimeraX.

The *B. burgdorferi* HMGR structures together suggest a catalytic mechanism similar to what has been proposed for the bacterial HMGRs^49^ (**Fig. S6**). In the *B. burgdorferi* enzyme, Glu 85, Lys 262, and His 375 on the CTD are positioned sufficiently close to the substrate for catalysis. The carboxylate of Asp 278 is within 2.58 Å of the amine of Lys 268, providing the proper positioning necessary for the Lys to also participate in H-bonding with Glu 85 and the substrate. Crucial to HMGRs is this microenvironment that keeps Glu 85 protonated (even at physiological pHs), allowing it to donate a proton after hydride transfer to the thioester. The collapse of the Glu 85 anion leads to CoA-SH release (after it receives a proton from His 375) and the formation of the mevaldehyde intermediate. In our structure, the CTD that houses His 375 appears to be in a “partially closed” state with the residue modeled 4.89 Å away from HMG-CoA (**Fig. 5B**). This is unsurprising, as our structure does not have cofactor present and would therefore not be expected to have proper orientation of the CTD. Following CoA-SH release, the mevaldehyde intermediate is then reduced by another equivalent of NAD(P)H, with Glu 85 again donating a proton to form the mevalonate product.

Considering the similarity of the proposed catalytic mechanisms, the conservation of key active site residues between HMGRs, but the lack of broad-spectrum inhibition by statins and the phenyl sulfonamides, we speculated that there might be unrecognized or unappreciated differences in the active site architectures of HMGRs that prevent these inhibitors from exhibiting cross-species reactivity. If true, these differences could be leveraged to design or discover species-selective HMGR modulators. This speculation is supported by data showing the evolution of these enzymes: HMGRs are incredibly diverse outside certain regions of their active sites (**Fig. 6C**). Many of the surface residues at locations distal from the active site show substantial differences in their amino acid composition. While there is much greater conservation at the active site, there are locations where the sequence divergence is notable. One example is the region close to the adenosine ring of the substrate – the large sequence divergence here suggests that the enzymes all have differences in their positioning of the HMG-CoA substrate. Given that statins are mimics of HMG-CoA, it is plausible that slight differences in the architecture of the substrate binding cavities impacts inhibitor engagement. It is also possible that the conformational change to properly position the catalytic residues of the C-terminal domains also contribute to variance in the architectures of the active sites.

## Discussion

Lyme disease is a prevalent vector-borne disease. While treatable with existing antibiotics, its rising incidence increases the susceptibility of the causative agent to antimicrobial resistance. Thus, new strategies to curtail the pathogen are needed. Because *B. burgdorferi* outsources many key aspects of its metabolism to infected hosts, we hypothesized that it might be vulnerable to disruptions of primary metabolic pathways. In this work, we used direct methods to demonstrate that the mevalonate pathway is essential for the proliferation of *B. burgdorferi* in vitro, and that its essentiality is directly related to de novo synthesis of peptidoglycan. CRISPR interference targeting the rate-limiting HMGR caused severe cell morphological defects that were largely rescued with IPP supplementation. While IPP is an important precursor to isoprenoids that decorate proteins, it is also necessary to construct peptidoglycan lipid carriers. Our investigations suggest that HMGR depletion impacts the latter function but does not rule out the disruption of other important cellular pathways that use isoprenoids.

In-depth biochemical characterization of the HMGR from *B. burgdorferi* identified that the enzyme that displays a high affinity for HMG-CoA with an apparent *K_M_* similar to what the human enzyme displays for substrate binding but that the enzyme is not disabled by potent active site inhibitors of the human ortholog. Statins that are potent inhibitors of human HMGR and two phenyl sulfonamides that inhibit the *E. faecalis* HMGR showed no effect on the activity of the *B. burgdorferi* enzyme, indicating that, despite performing the same reaction using conserved catalytic residues, HMGR enzymes possess important differences that influence susceptibility to small molecule inhibitors. Crystal structures of the *B. burgdorferi* enzyme bound to various ligands provided unprecedented insights into the ligand-binding site and support the notion that the subtle structural differences related to substrate positioning likely impact enzyme function. These structures revealed why, despite the suggestions that statins be used as antibiotic adjuvants, these drugs are poor antimicrobial agents: the HMGR active sites are distinct and lack many of the key polar interactions that stabilize statins in the human HMGR active site. Our modeling suggests that as many as six hydrogen bonding partners are absent in the *B. burgdorferi* HMGR that together could lower the equilibrium affinity of statins by 400-fold. It also appears that this active site distinction is not limited to the *B. burgdorferi* HMGR, as the structural mapping of sequence diversity reveals substantial active site differences at positions occupied by residues not involved in catalysis. This remarkable divergence may explain the lack of potent broad-acting HMGR inhibitors and could be used to guide the discovery of species-selective active site modulators.

One striking finding from this study was the discovery that the *B. burgdorferi* HMGR and homologs possess a cofactor motif different from those established for enzymes preferring NADH and NADPH. Such a finding was only made possible by access to thousands of HMGR sequences deposited in public repositories. Sequence and structural analyses of these proteins bearing the non-canonical cofactor motif revealed remarkable conservation at positions previously thought to be more variant. Moreover, because the sequences the divergent motif have elements of both NADH and NADPH motifs, we suspected that the enzymes bearing them could function with either nicotinamide cofactor. Indeed, this is the case, as the majority of homologs investigated have catalytic efficiency ratios for the two cofactors that are lower than observed for many oxidoreductases and certainly for most HMGRs. While the exact reason for this promiscuity is not known, it might allow greater adaptability for metabolic regulation. One possible explanation for the evolutionary acquired trait of cofactor promiscuity can be explained by the complex lifecycle of the bacteria in which the enzymes operate. *B. burgdorferi* has a complex lifecycle as it alternates between tick and vertebrate hosts, where the environments are metabolically distinct^9^. These niches could have variations in cofactor availability and redox balance, which could induce selective pressure on tightly regulated metabolic enzymes. Therefore, having an enzyme that can utilize both cofactors may serve as an adaptive strategy to regulate metabolic flux through the MVA pathway under fluctuating metabolite availability. Future work is being conducted to investigate this evolutionary link of MVA metabolic adaptation in host-dependent pathogens.

Beyond any purported physiological relevance, the nicotinamide cofactor promiscuity could be exploited to enable the facile engineering efficient metabolic pathways. The ability to alter cofactor specificity in oxidoreductases has substantially impacted efforts to engineer microbes for the bioproduction of valued natural products. For enzymes such as HMGRs that are directly relevant to the biosynthesis of isoprenoids (including ginkolides and Taxol), attempts to engineer organisms to produce these high value natural products are hindered by dysregulated carbon metabolism, with the engineered pathways tending to syphon key resources needed to sustain host metabolism. Published work has shown that a more even balance of redox cofactors can better regulate pathway engineering, but this is difficult to achieve given the remarkable cofactor selectivity of oxidoreductases^37, 50, 51^. By identifying enzymes that are already exhibit cofactor promiscuity, this work sets the stage to better explore the tuning and altering of cofactor selectivity, with aims to ultimately employ these enzymes in pathway engineering. There is only one other HMGR homolog (from A. *fulgidus*^29^) reported to display cofactor plasticity in the range discovered for enzymes in the undefined clades. While it remains unclear what drives the promiscuity in that case, we posit that an understanding of the inherent active site plasticity of HMGRs will be critical for their eventual deployment as biocatalysts tolerant to different redox cofactor pools.

In conclusion, this work establishes the essentiality of the mevalonate pathway in the metabolically minimal pathogen *B. burgdorferi* and the cofactor promiscuity of the rate limiting HMGR enzyme. This pathway represents the second example of a primary metabolic pathway that is a point of vulnerability for the pathogen, with the first being lactate dehydrogenase, the enzyme responsible for synthesizing lactate from pyruvate^52^. Beyond *B. burgdorferi*, this work reveals that, despite decades of investigation into HMGRs, key elements of their mechanism remain unknown. We posit that this work employing direct visualization methods to confirm the necessity of HMGR for *de novo* peptidoglycan synthesis will elevate these enzymes as targets for the development of novel antimicrobials while illuminating their potential for synthetic biology applications.

## Methods

### Borrelia burgdorferi strain and growth conditions

All *B. burgdorferi* strains used in this study are derived from a B31 K2 strain^53^. All *B. burgdorferi* cultures were grown in complete Barbour-Stoenner-Kelly media^54^ (BSK-II) containing 50 g/L BSA (Millipore 81-003), 9.7 g/L CMRL-1066 powder (USBiological C5900-01), 5 g/L neopeptone (BD 211681), 6 g/L HEPES Acid (Millipore 391338), 5 g/l D-glucose (Sigma G7021), 2 g/L yeastolate (BD 255772), 0.7 g/L sodium citrate (Sigma C7254), 0.8 g/L sodium pyruvate (Sigma P5280), 2.2 g/L sodium bicarbonate (Sigma S5761), and 0.4 g/L *N*-acetyl-glucosamine (Sigma A3286). All these components were mixed, the pH was leveled to 7.6, then the medium was brought to 1 L and filter sterilized. Following, 60 mL/L of heat-inactivated rabbit serum (Gibco 161120099) was added to the medium. The cultures were kept below 5×10^7^ cells/mL. *B. burgdorferi* cells were hand-counted by use of disposable hematocytomers (InCyto DHC-N101) under dark-field microscopy, as described previously^34^. Relevant antibiotic concentrations were 100 µg/mL streptomycin and 200 µg/mL kanamycin. For induction, 100 µM IPTG (Sigma I6758) was used. For supplementation where relevant, 2.5 mM mevalonate (Sigma 50838) or IPP (Sigma 39784) were used. For resuspending cell pellets for microscopy, complete BSK-II medium without phenol was used. This medium contains all components used in regular BSK-II but instead uses CMRL-1077 without phenol red (USBiological C5900-03A).

All *E. coli* cultures were grown either in pre-mixed lysogeny broth (LB) containing 1% tryptone, 0.5% yeast extract, and 5 g sodium chloride per liter (Fisher BP1426-2) and cultured at 30 °C shaking 220 rpm or plated on LB plates containing 1.5% agar (Sigma A5306). For recovery after heat-shock transformation, cells were suspended in SOC medium containing 2% tryptone, 0.5% yeast extract, 10 mM NaCl, 2.5 mM MgSO_4_, and 20 mM glucose (MP Biomedicals 113031012). For growing cultures to harvest plasmids using midiprep kits, cells were grown in super broth containing 12 g soy hydrolysate, 24 g yeast extract, 11.4 g K_2_HPO_4_ and 1.7 g KH_2_PO_4_ per liter (Fisher BD 212485). *E. coli* antibiotic concentrations were 50 µg/mL for streptomycin.

### BSK-II agarose pad preparation

For time-lapse microscopy, cells were spotted on agarose pads containing BSK-II medium to promote growth. Due to heat-labile components in BSK-II, such as BSA or rabbit serum, the medium was prepared as a separate 3× concentrate and added to a cooled agarose solution. This 3× BSK-II concentrate contained 100 g/L BSA (this approaches the solubility limit of BSA, Millipore 81-003),29.1 g/L CMRL-1066 without phenol red (to reduce autofluorescence in microscopy, US Biological C5900-03A), 15 g/L neopeptone (BD 211681), 18 g/L HEPES acid (Millipore 391338), 15 g/L D-glucose (Sigma G7021), 6 g/L yeastolate (BD 255772), 2.1 g/L sodium citrate (Sigma C7254), 2.4 g/L sodium pyruvate (Sigma P5280), 6.6 g/L sodium bicarbonate (Sigma S5761) and 1.2 g/L *N*-acetyl-glucosamine (Sigma A3286). All components were mixed, pH was brought to 7.6, then the medium brought to 500 mL and filter-sterilized in a 0.2 µm filter sterile disposable bottle (Fischer Scientific 09-741-02) for five hours due to the high lipid and protein content in the medium. After filtration, 45 mL/L of heat-inactivated rabbit serum (Gibco 161120099) was added to the medium. The 3× BSK-II solution was frozen in 50 mL portions, from which one could be thawed and aliquoted in 1 mL stocks and frozen again, ready for experimental use.

Seaplaque GTG Agarose (Lonza 50111) was mixed in milliQ H_2_O to a concentration of 4.5%, vortexing thoroughly to mix until the agarose was fully suspended in a 1.7 ml Eppendorf tube. The solution was polymerized on a heat block set to 80°C for ∼1 hour, vortexing once every 10 min. Once ready, the heat block was lowered to 45°C. A 1 mL 3× BSK-II aliquot was then thawed at 45°C on the same heat block. Once both agarose and BSK-II were equilibrated to temperature, 3× BSK-II was added to bring the agarose to a final 3% concentration and BSK-II to 1× concentration, vortexing intermittently over 5 min to homogenize the mixture. Three percent agarose was used to reduce the mobility of *B. burgdorferi* over the course of the timelapse. One hundred microliter agarose pads were poured on a slide. They were set to gel for 30 min to reduce their moisture content before spotting cells to reduce *B. burgdorferi* motility. The agarose pad was sliced in half. To one half, 0.5 µL of 10 mM IPTG was added; the other half received 0.5 µL of milliQ H_2_O. These slices were allowed to dry for another ∼3 min until the liquid had mostly evaporated.

### HADA staining

Cultures of the *hmgr* CRISPRi strain (CJW_Bb662) at density of ∼10^7^ cells/mL were diluted to 1×10^6^ cells/mL in fresh BSK-II containing 100 µg/mL streptomycin and supplemented with nothing, 100 µM IPTG, 100 µM IPTG + 2.5 mM mevalonate, or 100 µM IPTG + 2.5 mM IPP. The cell density of these cultures was determined once daily. By day 2 post-supplementation, 500 µL of culture was taken from each culture and mixed with 1 µL of 50 mM HADA. They were incubated in the dark at 34 °C for 1 hour after which the samples were washed twice with 500 µL fresh BSK-II, centrifuged at 9000 × *g* for 5 min at room temperature. The final pellets were resuspended with 100 µL of BSK-II without phenol red to reduce autofluorescence on the microscope. An aliquot (1µL) of cell suspension was spotted on a fresh 2% PBS agarose pad, allowed to dry for 1 min, covered with a 0.15 mm coverslip, and sealed with VALAP, a 1:1:1 mixture of Vaseline petroleum jelly (Amazon ASIN B07MD6HJT4), lanolin (Spectrum # LA109) and paraffin wax (Fisher # 18-607-738). This sample was taken to the microscope for imaging.

### Snapshot microscopy setup

For microscopy snapshots, 5 µL of CJW_Bb662 culture supplemented with nothing, 100 µM IPTG, 100 µM IPTG + 0.5 mM IPP, or 100 µM IPTG + 2 mM IPP were spotted on 100 µL 2% agarose PBS pads^36, 55^. Samples were allowed to dry for ∼5 min before being sandwiched with a 0.15 mm coverslip and sealed with VALAP. Samples were imaged by microscopy immediately.

### Time-lapse microscopy setup

Cultures of the *hmgR* CRISPRi strain (CJW_Bb662) at a density of ∼10^7^ cells/mL were diluted to 1×10^6^ cells/mL in fresh BSK-II containing either 100 µg/mL streptomycin only (no induction) or with 100 µM IPTG (induction of CRISPRi). On day 2 post supplementation, the cultures were counted for density and then prepped for time-lapse microscopy. During the 30-min agarose pad gelling described above, 1- and 3-mL aliquots were taken from non-induced and IPTG-induced cultures, respectively, to account for differences in cell density due to HMGR depletion, then all tubes were spun at 9000 × *g* for 5 min. The supernatants were removed. From there, the “non-induced” aliquot was resuspended with 150 µL of 1× BSK-II without phenol red (prepared from the leftover thawed 3× BSK-II aliquot in the agarose pad preparation described above). The pellets from the “induced” samples were resuspended by adding 75 µL to one pellet, aspirating to mix, then transferring that volume to each pellet and pooling their cells for one aliquot. A cell suspension (0.5 µL) was spotted on their respective agarose pad (with or without IPTG), allowed to dry for ∼5 min, sandwiched with a 0.15 mm coverslip, and finally sealed with VALAP. This sample was taken to the microscope for imaging.

### Microscopy

For counting *B. burgdorferi* cells in disposable hematocytometers by dark-field microscopy, a Nikon Eclipse E600 microscope equipped with a 40× 0.55 NA Ph2 phase contrast air objective and a darkfield condenser lens was used. For fluorescence microscopy, specimens were imaged using a Nikon Ti inverted microscope equipped with a 100× 1.5 NA Ph3 phase contrast oil objective, Hamamatsu Orca-Flash4.0 V2 CMOS camera, and either a Sola LE light source (phase contrast) or a Spectra X Light engine (fluorescence, Lumencor). A “DAPI” filter set was used to acquire fluorescent images (ET395/25 ex, dichroic T425lpxr, ET460/50 em). The software NIS-Elements-AR was used to interface with the microscope and to control it. Time-lapse images were acquired once every 5 min. At every region of interest, the microscope auto-focused within a range of ± 5 µm using a step size of 0.5 µm. An Okolab incubator was used to maintain the microscope enclosure at 37°C.

### Image analysis

FIJI^56, 57^ was used to initially check and prepare images. Otherwise, image analysis was completed in Python for the remainder of the work using a previously published pipeline^58^. This pipeline uses the packages numpy^59^, scipy^60^, scikit-image^61^, seaborn^62^, and pandas^63^. Additionally, to facilitate analysis, large ND2 files were loaded and unpacked using the function NDtwoPy^64^. Phase-contrast images were binarized into masks using an adaptive threshold from the Python package OpenCV (*cv2.adaptiveThreshold*) and objects less than 500 square pixels were excluded. Surviving masks were skeletonized and used to measure the instantaneous width along the mask. Cells at least 5-µm long were retained. All surviving binary objects were manually curated as masks in FIJI, assigned an identity (ID), and then archived in a Pandas data frame for fluorescence measurements. Cells were considered “bulging” if their cell width exceeded 0.7 µm.

To measure the fluorescence of each cell object, images were first background-subtracted using a modified background subtraction algorithm^65^. Phase-contrast images were segmented using OpenCV’s adaptive threshold as above (*cv2.adaptiveThreshold*) to remove potential fluorescing objects. Masks were dilated through ten iterations of Scipy’s *binary_dilation* function^60^. These dilated objects were removed from the fluorescent image. Surviving pixels were binned and averaged using a rolling window across the *x* and *y* dimensions and smoothed with a Gaussian kernel to obtain an estimated “background-only” image. Cell masks were used to measure the mean intensity (integrated intensity per unit pixel area, or signal concentration) from every cell in the background-subtracted images. Histograms of single cell intensities were fit to a two-population Gaussian distribution (Eq. 1 below), where *P* is the percent contribution for each population while *µ* and *σ* are the mean and standard deviation of each Gaussian distribution, respectively.

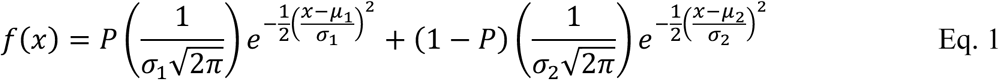

As described above, time-lapse images were checked initially using FIJI. Large ND2 files were loaded and unpacked using the NDtwoPy package^64^. and the movies were drift-corrected using the MATLAB package Supersegger^66^. Movies were then prepared in Python using scikit-image’s function *imread*^61^, frames were individually cropped, plotted, and labeled using Matplotlib. These individual slices were loaded into FIJI, where they were saved as “.AVI” movie files.

### Plasmid construction

For pBbdCas9S(RBSmut)-P_syn_-sgRNA^BB0685^, oligo pair 5’-AGTGATTAAAAAGAAAATCTTCAT -3’ and 5’- AACATGAAGATTTTCTTTTTAATC -3’ were phosphorylated using T4 PNK (NEB M0201S) and ligated to SapI-digested pBbdCas9S(RBSmut)-P_syn_-sgRNA500 using T7 DNA ligase (NEB M0318S). The resulting product was then transformed into chemically competent *E. coli* NEB C2992H (New England Biolabs). Clones were screened using whole plasmid sequencing. Plasmid was further verified by sequencing.

### Strain construction

For CJW_Bb662, plasmid pBbCas9S(RBSmut)_P_syn_-sgRNA^BB0685^ was electroporated into strain B31 K2 and plated on BSK-II agar plates supplemented with 100 µg/mL streptomycin. This isolated clone was confirmed to have the full B31 K2 plasmid complement by plasmid profiling.

The sequence of the insert coding for the gRNA targeting *bb0685* (*hmgr*) was verified by sequencing.

### Plasmid construction and mutagenesis

A codon-optimized *bb0685* with a sequence encoding a N terminal 6×-histidine tag cleavable via both TEV and thrombin was cloned into a pET28b plasmid at the NdeI and XhoI restriction sites. The plasmids were then transformed into Stellar cells (*E. coli* HST08), miniprepped, and confirmed through sanger sequencing (GeneWiz).

To create the MBP-*bb0685* fusion construct, a codon-optimized, the HMGR gene was cloned into a pMAL-C5× vector containing a N-terminal MBP that was cleavable via Factor Xa (NdeI and XhoI restriction sites).

The genes encoding the HMGRs from *L. jensenii* (A0A5N1IG17), *F. psychrophillium* (A0A076P6Q4), and *L. pneumophilia* (Q5ZTV6) were codon-optimized and sub-cloned into pET28a vectors with an N terminal 6×-histidine tag cleavable via the TEV protease.

### RNA extraction and qRT-PCR

RNA extraction and qRT-PCR were performed as previously described^67, 68^. Three replicates of CJW_Bb662 cells were grown to a density of 10^7^ cells/mL in fresh BSK-II, then incubated with 100 µM IPTG for 4 h to induce CRISPRi. Cells were pelleted by centrifuging at 4,300 × *g* for 10 min after which the supernatant was removed. Cells were resuspended by pipetting in 400 µL HN buffer (50 mM NaCl, 10 mM HEPES, pH 8.0) before adding 1 mL of RNAprotect Bacteria Reagent (Qiagen 76506). Samples were vortexed to ensure homogeneity, followed by an incubation at room temperature for 5 min. Cells were then centrifuged at 10,000 × *g* for 10 min, after which the supernatant was removed. Cell pellets were frozen at −80°C until RNA extraction was performed using protocol 4 in the RNAprotect bacterial reagent handbook, which includes enzymatic cell lysis and proteinase K digestion. RNA purification was performed using RNeasy Mini Kit (Qiagen 74014) as described in protocol 7 of the RNAprotect bacterial reagent handbook. DNase digestion was then performed using the TURBO DNA-free kit (Thermo Scientific AM1907) following the rigorous DNase treatment protocol. Purified RNA was stored at −80°C.

RNA was quantified using the KAPA SYBR FAST qPCR Mastermix (2×) Universal kit (Roche KK4650). Reactions were performed in duplicate or triplicate, using10 ng of total RNA per reaction. Triplicate reactions were also performed without the addition of the reverse transcriptase. The primers used for amplification were 5’- TTCAATCAGGTAACGGCACA-3’ and 5’-GACGCTTGAGACCCTGAAAG-3’ for the control *flaB*, 5’-GCATTTGCTACCGGGTAAAG-3’ and 5’-CGCTCCTCTTCGTAAAAACC-3’ for *hmgr*, 5’-AGGCCCCCAAGTAAAGTTTC-3’ and 5’-CCATTCTAAGTCACACCCAACC-3’ for *mvaD*, 5’-GCAATTGGAGAGGCAATAGG-3’ and 5’-CGGCTCCCAAAGCTTTAATC-3’ for *pmk*, and 5’-GCGTTTGGGGTTGTCTAATG-3’ and 5’-CAGCGCCACTTAACTTACCAG-3’ for *mvk*.

Using a Bio-Rad CFX Connect Real-time system, the following protocol was performed: reverse transcription (5 min at 42°C), enzyme activation (3 min at 95°C), 40 cycles of annealing, extension, and data acquisition (3 s at 95°C, and 20 s at 60°C with fluorescence acquisition in the SYBR scan mode), and a melt curve analysis (55°C to 95°C in 0.5°C increments). The *hmgr, mvaD, pmk,* and *mvk* transcript levels under CRISPRi-inducing (+IPTG) condition were normalized to the level of *flaB* transcript, and then expressed as relative to the normalized transcript level under the non-inducing condition (-IPTG) using the 2^-ΔΔCT^ method^69^.

### Flow cytometry

CJW_Bb662 parent cultures (10^6^ cells/mL) were diluted to 5×10^4^ cells/mL in fresh BSK-II with 100 µg/mL streptomycin supplemented with nothing, 100 µM IPTG, 100 µM IPTG + 0.5 mM mevalonate, 100 µM IPTG + 2 mM MVA, 100 µM IPTG + 0.5 mM IPP, or 100 µM IPTG + 2 mM IPP. Conditions were performed in triplicate. These cultures were counted daily by flow cytometry using a ThermoFisher Attune CytPix Flow Cytometer. Cultures were diluted as needed in fresh BSK-II for accurate detection within the dynamic range of the detector. Samples (50 µL) were run at a rate of 25 µL/min. To prevent carryover, 150 µL of water was run between samples at a rate of 200 µL/min.

Flow cytometry data were analyzed using FlowJo (version 10.10). Gating thresholds were determined by comparison of BSK-II medium only, exponentially growing CJW_Bb662, and IPTG-induced CJW_Bb662. This methodology allows for variable morphological phenotypes. After gating, cell counts were multiplied by their dilution factor to determine the cell density (cells/mL).

### Plate reader absorbance assay

Based on previous protocols^34, 70^, K2 cells were diluted to 1×10^4^ cells/mL in fresh BSK-II and supplemented with 1% DMSO, 100 µg/mL streptomycin, 1% DMSO + 20 µM lovastatin, 1% DMSO + 50 µM lovastatin, or 1% DMSO + 800 µM atorvastatin. Conditions were performed in triplicate. After the setup was complete, day 0 absorbance values were taken every 10 nm from 350 to 650 nm to cover the absorbance peaks of phenol red using a BioTek Synergy2 plate reader. Samples were then incubated at 34°C and 5% CO_2_ for 7 days. On day 7, the samples were removed from the incubator and allowed to acclimate to room temperature for 24 h. Absorbance values were taken again on day 8. Absorbance measurements were taken at room temperature to prevent effects associated with temperature and dissolved CO_2_. The data were analyzed in Python using built-in pandas functionality. Absorbance values for days 0 and 8 were loaded as separate data frames, where the values from day 0 were subtracted from the values from day 8 to yield ΔAbsorbance.

### Protein expression and purification of *B. burgdorferi*, *L. jensenii*, *F. psychrophillium* and *L. pneumophilia*, and *L. jensenii* R167T variant HMGR

All plasmids were transformed into *E. coli* BL21 chemically competent cells (New England Biolabs, MA). Transformed cells were grown in autoinduction media (ZYM-5052) supplemented with 100 mg/L ampicillin at 37°C. Cells were cultured while shaking at 220 rpm until OD_600_ reached value between 0.6 – 0.8, at which point the temperature was shifted to 18 °C overnight. Cell cultures were then centrifuged at 8,000 × *g* for 30 min and the pellets frozen with liquid nitrogen and stored at −80°C.

Frozen cell pellets were resuspended in lysis buffer (25 mM HEPES pH 7.5, 200 mM NaCl, 5 mM β-mercapto ethanol or β-ME, 10% glycerol), 10 mg/L lysozyme, 10 mg/L DNAse, 1 M MgSO_4_, and 0.5 mg/L phenylmethylsulfonyl fluoride (PMSF, diluted in 50/50 ethanol/water). The suspension was lysed using Microfluidizer LM20 (Microfluidics) at 17,000 psi. After lysis, 0.45 mg/L PMSF was added to the lysate, and the cells were centrifuged at 22,000 × *g* for 30 min to clarify the lysate. The cleared lysate was loaded onto an amylose gravity-flow column (∼10 mL resin per 100 mL lysate) pre-equilibrated with lysis buffer. The column was washed with 10 column volumes (CV) of lysis buffer and eluted with 10 CV of elution buffer (25 mM HEPES pH 7.5, 200 mM NaCl, 5 mM β-ME, 10% glycerol, 10 mM maltose). Protein purity was assessed via SDS-PAGE analysis, and the fractions containing the fusion protein were concentrated at 4,000 rpm using a 10 kDa Amicon Ultra-15 Centrifugal Filter Unit (Tullagreen, Carringtwohill Co. UFC9010). Concentrated protein was then aliquoted into 1 mL fractions and flash frozen using liquid nitrogen and stored at −80 °C. 20 g of cell pellet yielded 40 mg/mL of protein. Following concentration, MBP was cleaved by the addition of 2% (w/w) Factor Xa protease (New England Biolabs P8010L) overnight at 4 °C; the reaction components were separated by analytical size exclusion chromatography. Purity of samples was again assessed via SDS-PAGE analysis and concentrated.

Each homolog was purified as described above; pure proteins were aliquoted into 1 mL fractions, flash frozen using liquid nitrogen, and stored at −80 °C. Twenty grams of cell pellet yielded 100 mg/mL of protein *L. jensenii* HMGR, 40 mg/mL of *F. psychrophilium* HMGR, and 10 mg/mL of *L. pneumophilia* HMGR.

### Analytical size exclusion chromatography

100-250 μL of protein sample at the indicated concentrations were injected into a 25 mL analytical Superdex 200 Increase column (10/300 GL; Cytiva 28990944) and separated at a flow rate of 0.250 mL/min. The absorbance at 280 nm was recorded, and 0.5 mL peak fractions were collected. Purity of the protein fractions was assessed through SDS-PAGE analysis.

### Kinetics characterization

All kinetic assays were performed by measuring the enzyme-catalyzed oxidation of NAD(P)H that is observed via the decrease in absorbance at 340 nm using a Tecan Spark plate reader and an extinction coefficient of 6200 M^-1^ cm^-1^ for NAD(P)H. Kinetics of cofactor specificity was analyzed in 100 μL reactions at 30 °C with the following reaction mixture: 25 mM HEPES (pH 7.5), 200 mM NaCl, 5 mM β-ME, 200 μM HMG-CoA, 3 – 330 μM NAD(P)H and 0.5 μM enzyme. For kinetics of HMG-CoA, reactions were carried out at 30 °C in a total volume of 100 μL with the following reaction mixture: 25 mM HEPES (pH 7.5), 200 mM NaCl, 5 mM β-ME, 400 μM NAD(P)H, 2 – 60 μM HMG-CoA, and 0.5 μM *B. burgdorferi* HMGR. Kinetics parameters were obtained by using nonlinear regression in Prism (GraphPad Software) to obtain initial velocities. Rates versus substrate/cofactor concentration was fit to the Michaelis-Menten equation and the Michaelis-Menten constant (*K_M_*) and the maximum velocity (*V*_max_) were obtained. Some curves were fit with the allosteric sigmoidal equation 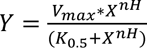 to obtain Hill coefficient (n_H_) and *K*_0.5_.

The activity *vs* pH profile of the HMGR from *B. burgdorferi* was determined as described above, but with the following reaction mixture: 0.05 M citric acid, 0.1 sodium phosphate dibasic, 200 mM sodium chloride, 5 mM β-ME, 150 μM HMG-CoA, 300 μM NAD(P)H, 1 μM *B. burgdorferi* HMGR, and adjusted pH from 4 – 9 using NaOH. Kinetics parameters were obtained by using nonlinear regression in Prism (GraphPad Software) to obtain initial velocities. Rates versus substrate/cofactor concentrations were fit to the Michaelis-Menten equation to obtain values of *K_M_* and *V*_max_.

### 8-anilino-1-napthalenesulfonic (ANS) acid fluorescence

Assays were performed in 25 mM HEPES (pH 7.5), 200 mM NaCl, 5 mM β-ME, 0.25 mg/mL *B. burgdorferi* HMGR, 30 μM ANS, 12 – 330 μM NAD^+^/HMG-CoA, in a final reaction volume of 50 μL. Substrates were titrated into the samples and incubated for 5 minutes in the dark. Each reaction was excited at 372 nm, with emission monitored and recorded from 400 – 600 nm at 25 °C in a Tecan Spark plate reader. Emission intensities at 470 were collected and normalized to the control.

### Crystallization of HMGR from *B. burgdorferi*

Sparse-matrix crystallization screens were performed with purified protein at ∼5.6 mg/mL. Using the sitting-drop vapor diffusion method, all crystals were obtained by mixing 0.5 μL of protein and 0.5 μL of precipitant solution at room temperature. Crystals of *B. burgdorferi* HMGR with 1 mM HMG-CoA first appeared in the MCSG1-C12 screen, which contains 0.1 M Bis-Tris HCl, pH 6.5, and 25% (w/v) PEG 3350, after 2 days. Crystals were cryoprotected in paraffin oil, flash-frozen, and stored in liquid nitrogen.

Optimization trays were performed with 0.1 M Bis-Tris HCl and varied pH from 5.5 to 7, and 19 to 29% (w/v) PEG 3350. Using the sitting-drop vapor diffusion method, all crystals were obtained by mixing 1 μL of protein and 1 μL of precipitant solution at room temperature. Crystals with 1 mM HMG-CoA, 1 mM NAD^+^, and 1 mM HMG-CoA/1 mM NAD^+^ appeared after 2 days. Crystals were cryoprotected in precipitant solution supplemented with 29% glycerol.

### Data collection, processing, and structure determination

All the diffraction data were acquired at 100 K at the Stanford Synchrotron Radiation Lightsource (SSRL, Stanford, CA, USA) on BL12-1 and BL12-2. The diffraction data were indexed and processed using XDS^41, 42^ and HKL3000 packages^43^. All structures were phased using Phenix Phaser with the AlphaFold2 multimer predicted structures of *B. burgdorferi* (AF-O51628-F1-v4) and *L. jensenii* (A0A5N1IG17) HMGRs. The structures were refined in Phenix Refine, and model building was performed in Coot. The statistics of data collection and refinement are listed in Tables 2 and 4. All structural figures were prepared using USCF ChimeraX^44^ and Adobe Illustrator.

### Phylogenetic analysis

Rooted maximum likelihood phylogenetic tree. The tree contains 210 sequences from the 14 largest clusters from the SSN with the eukaryotic HMGRs serving as an outgroup. Sequences were aligned with the MAFFT software (https://www.ebi.ac.uk/Tools/msa/mafft/). The maximum-likelihood phylogenetic rooted tree was subsequently computed with the IQ-tree software employing the LG+R8 model.

### Sequence Similarity Network (SSN) and Genome Neighborhood Diagram (GND)

Protein sequence similarity network (SSN) of bacterial HMGRs depicting size and functional clustering. The SSN was generated via the Enzyme Function Initiatives-Enzyme Similarity tool (EFI-EST, https://efi.igb.illinois.edu/efi-est/) and visualized in Cytoscape^71^ with an “organic” layout. The SSN was generated by employing a single sequence blast of *B. burgdorferi* HMGR and tailored so that the nodes represent sequences with 100% identity, an e-value of 5, and an alignment score of 125. There are 7,116 unique protein sequences. Genome neighborhood clustering of mevalonate pathway in *B. burgdorferi* strain B31. Genomic neighborhoods were generated via the EFI-Genome Neighborhood Tool with a reading frame of 10 and a minimal co-occurrence of 20%.

### Data and code availability

Raw microscopy images are available in the BioImage Archive^72^ under the accession number S-BIAD2681. Code for single-cell analysis and absorbance assays can be downloaded from the Jacobs-Wagner lab Github site: (<u>github.com/JacobsWagnerLab/published/tree/master/Paddy_et_al_Dassama_2026</u>).

Crystallographic data has been deposited in the Protein Data Bank^73^ and can be accessed with the following codes: 9ZW6, 9ZW7, 9ZW8,9ZW9.

## Acknowledgements

This work was supported in part by the National Institutes of Health grant R35GM150910 (to L.M.K.D.), the Howard Hughes Medical Institute Emerging Pathogens Initiative (to C. J-W. and L.M.K.D.), and the Innovative Medicines Accelerator. I.A.P. was supported by a Sarafan ChEM-H CBI Lipschultz Fellowship, is the recipient of an NSF GRFP, and is an HHMI Gilliam Fellow.

M.T.S. was supported by a Stanford DARE fellowship. C.J-W. is an HHMI investigator. L.M.K.D. was additionally supported by a Terman Fellowship from Stanford University and a MAC3 Impact Philanthropies Faculty Fellowship at the Sarafan ChEM-H Institute. Crystallography data was acquired at the Stanford Synchrotron Radiation Lightsource (SSRL). Use of SSRL and SLAC National Accelerator Laboratory is supported by the U.S. Department of Energy, Office of Science, Office of Basic Energy Sciences under Contract No. DE-AC02-76SF00515. The SSRL Structural Molecular Biology Program is supported by the DOE Office of Biological and Environmental Research, and by the National Institutes of Health, National Institute of General Medical Sciences (P30GM133894). The contents of this publication are solely the responsibility of the authors and do not necessarily represent the official views of NIGMS or NIH. The authors thank Dr. Daniel Fernandez at the Sarafan ChEM-H Nucleus for crystallographic data acquisition and are grateful to all members of the Dassama and Jacobs-Wagner groups for thoughtful discussions and insights.

## Movie legends

**Movie S1: Video of *B. burgdorferi* cells (control) growing and dividing on a BSK-II agarose pad.** Time-lapse phase-contrast microscopy of representative uninduced CJW_Bb662 cells.

**Movie S2: Video of *B. burgdorferi* cells undergoing HMGR depletion on a BSK-II agarose pad.** Time-lapse phase-contrast microscopy of representative CJW_Bb662 cells experiencing HMGR depletion on a BSK-II agarose pad containing the CRISPRi inducer IPTG.

## Supporting Information for

### Supporting figures

**Fig. S1.**
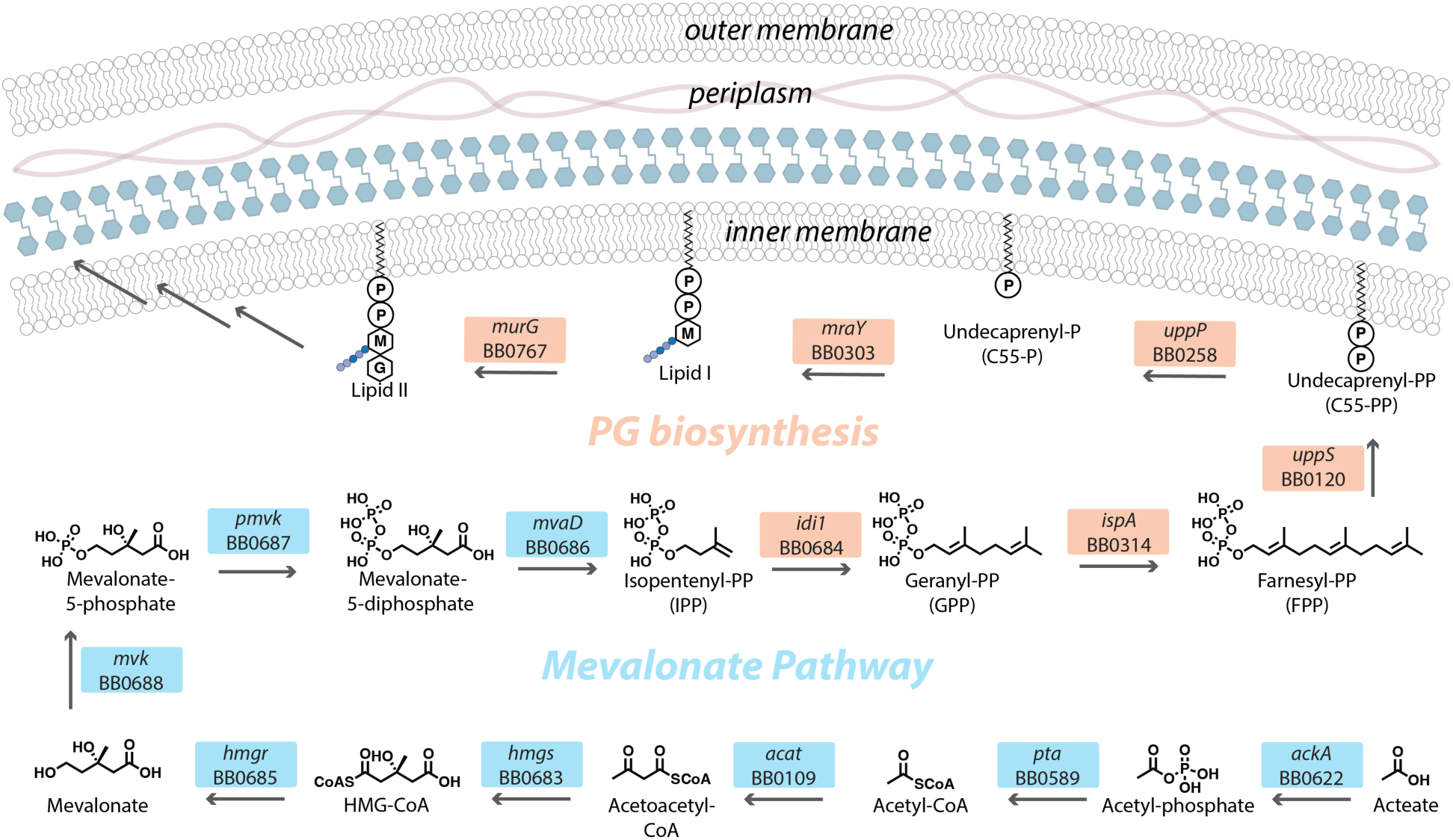
*B. burgdorferi* B31 mevalonate and peptidoglycan (PG) biosynthesis pathways. Enzymes in the mevalonate and PG biosynthesis pathways from *B. burgdorferi* B31 identified by their locus tags and gene names. Also shown are the chemical structures of key molecules that they act on.

**Fig. S2.**
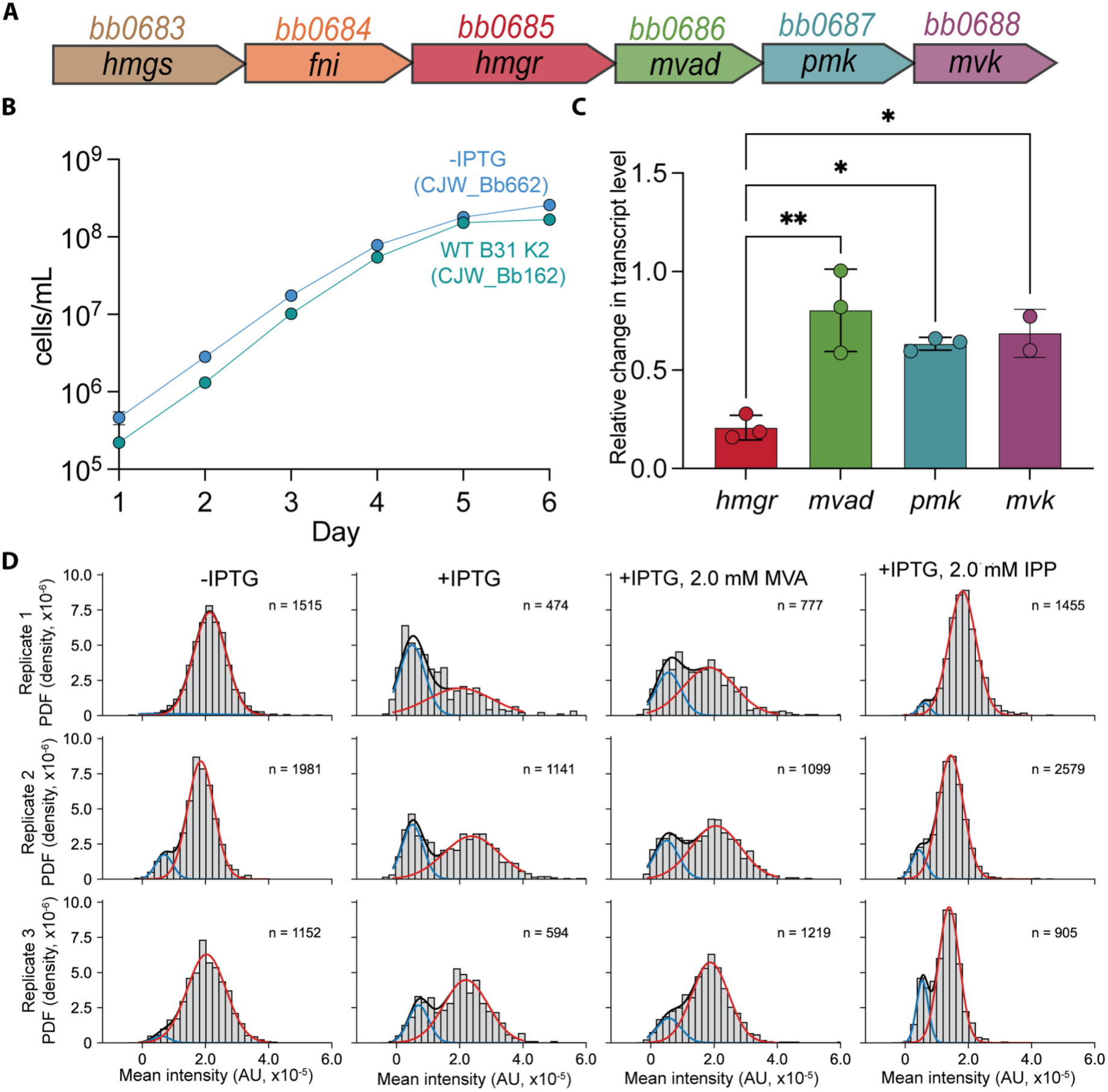
Effects of CRISPR-interference of *hmgr* and HADA labeling statistics. **A**) Genome neighborhood diagram of the MVA pathway in *B. burgdorferi* strain B31. **B**) Growth curves of CJW_Bb662 in comparison to wild-type strain in the absence of IPTG. **C**) Plot showing the *fla*B-normalized transcript levels of the indicated gene under IPTG-inducing conditions relative to the noninducing (-IPTG) condition. CJW_Bb662 cultures grown in BSK-II to 1 x 10^7^ cells/mL were split and either left untreated (-IPTG) or induced with 100 µM IPTG for 4 hours to trigger CRISPRi of *hmgr*. RNA was subsequently extracted from the samples and analyzed by qRT-PCR. Each dot represents a biological replicate. A one-way ANOVA with post-hoc Tukey test showed that the expression of *bb0686*, *bb0687*, *bb0688* is statistically different from that of *bb0685* (*hmgr*) (p < 0.05) but not statistically different from each other (p > 0.78). **D**) Distributions of single-cell HADA intensities for individual biological replicates (related to Fig. 1G-J) across the indicated conditions. The cell number (n) for each Gaussian-fitted population is provided. The black lines represent the fit for the two populations whereas the red and blue lines show the contribution of each population.

**Fig. S3.**
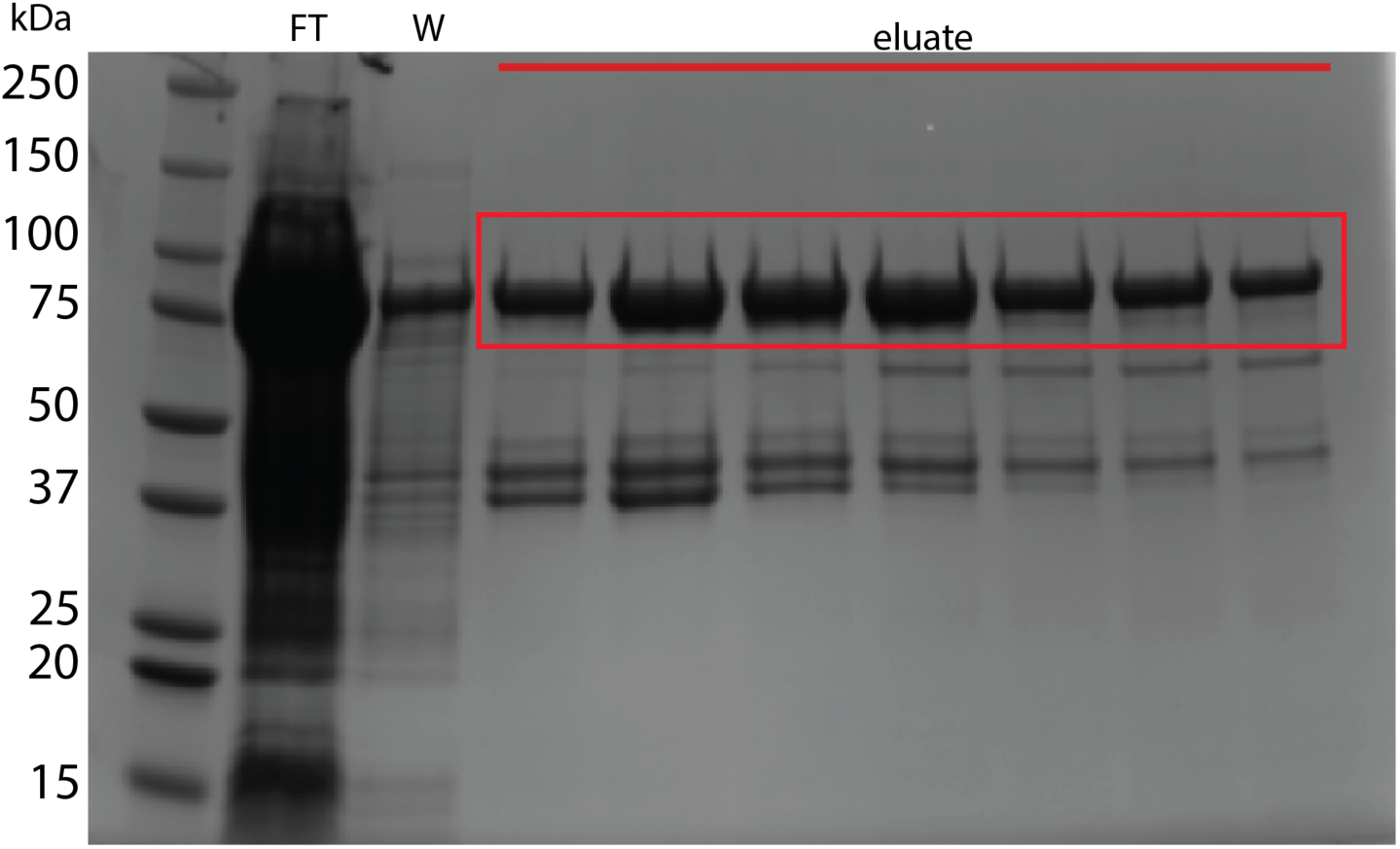
Purification of *B. burgdorferi* HMGR. SDS-PAGE gel of MBP-HMGR following dextrin affinity chromatography. Molecular weight of the MBP fusion is ∼91 kDa (red rectangle). FT: flow through; W: column wash.

**Fig. S4.**
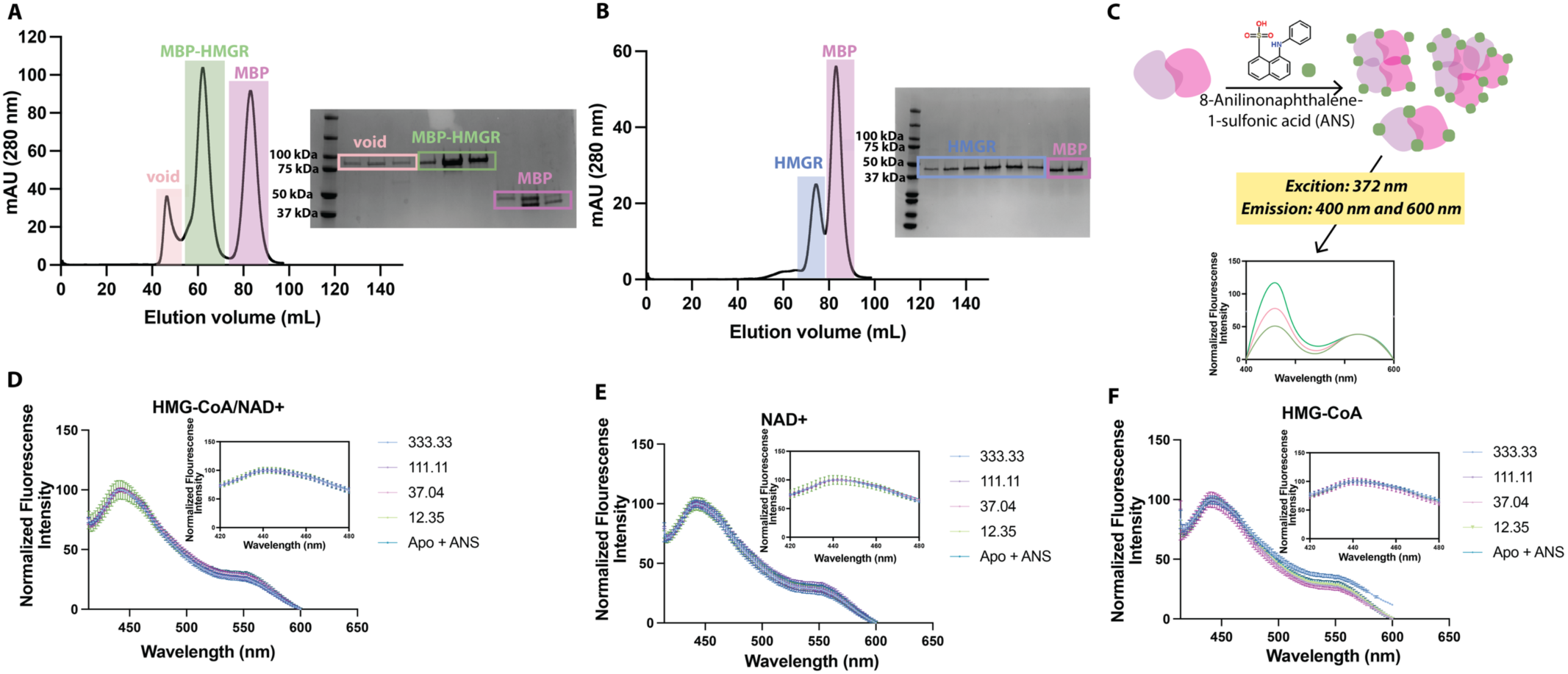
Biochemical analysis of *B. burgdorferi* HMGR. **A, B)** Size-exclusion chromatography of *B. burgdorferi* MBP-HMGR fusion post amylose purification (**A**) and following Factor Xa cleavage of MBP (**B**). **C**) Schematic of ANS (1-anilino-8-naphthalene sulfonate) fluorescence assay to detect oligomerization. **D-F**) Fluorescence *vs* wavelength spectra with ANS probe and HMGR from *B. burgdorferi* with the addition of HMG-CoA, HMG-CoA/NAD^+^, and NAD^+^, respectively.

**Fig. S5.**
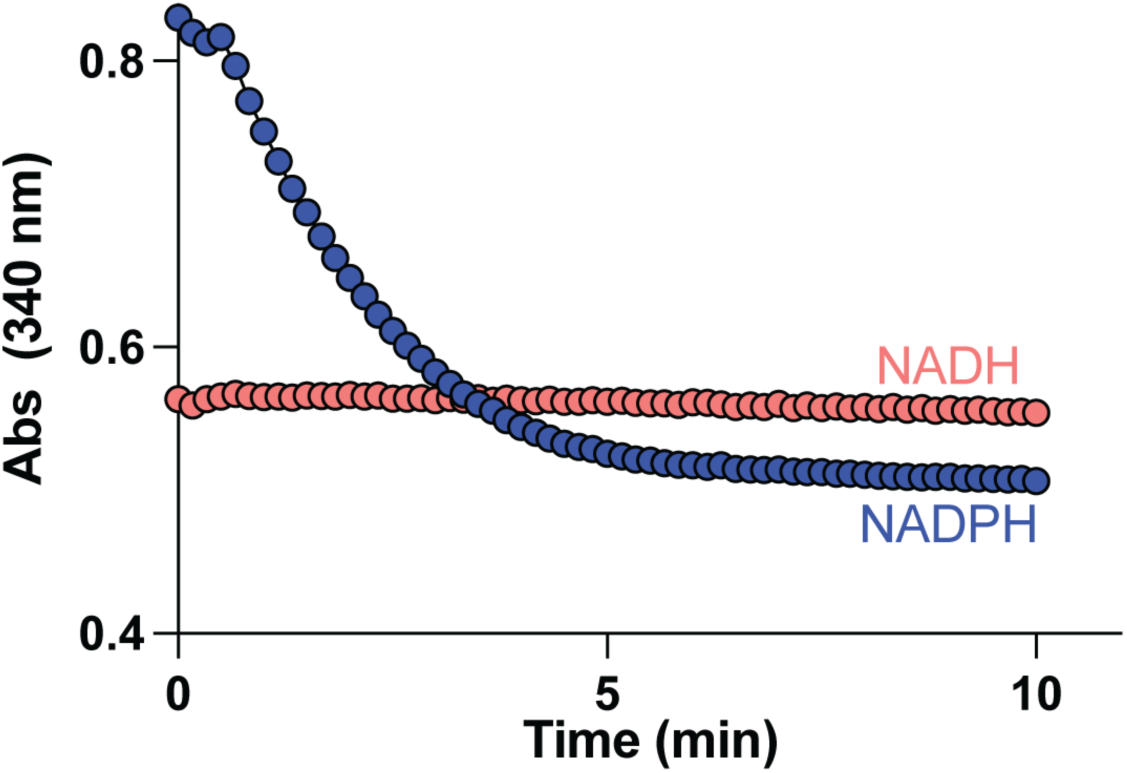
Absorbance plot of *H. sapiens* HMGR with both cofactors. Absorbance plot over time of the *Homo sapiens* HMGR reaction monitoring the oxidation of NADH (red) or NADPH (blue). Assay conditions were 10 nM enzyme, 125 mM HMG-CoA, 250 NADPH or NADH.

**Fig. S6.**
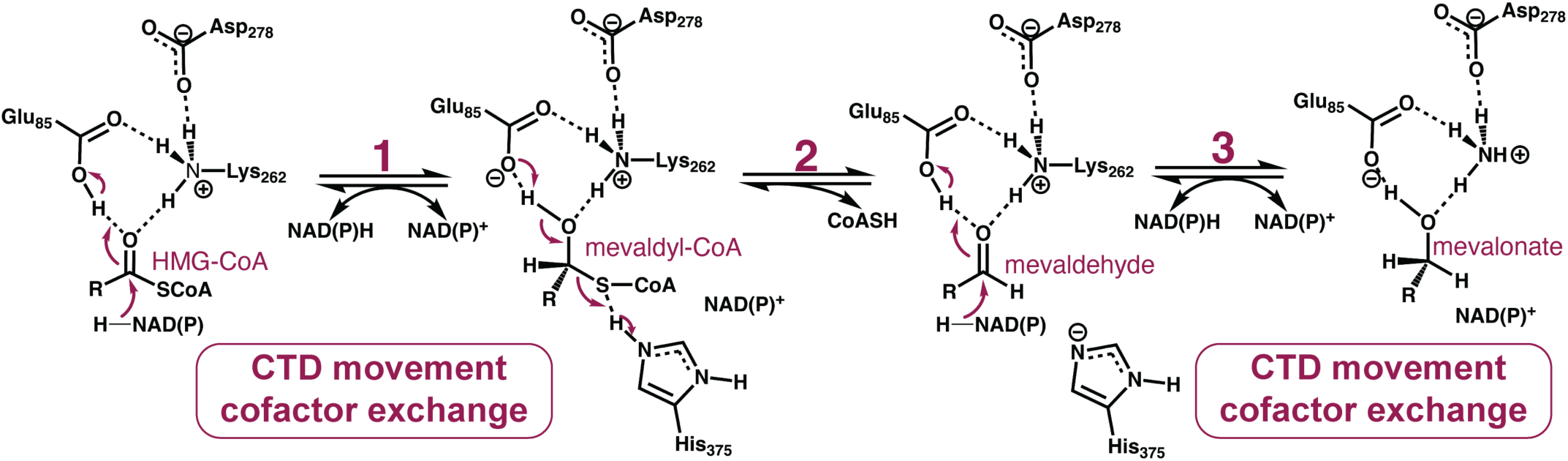
Bacterial HMGR reaction mechanism. Abbreviated proposed reaction mechanism of HMGR from *B. burgdorferi*.

**Fig. S7.**
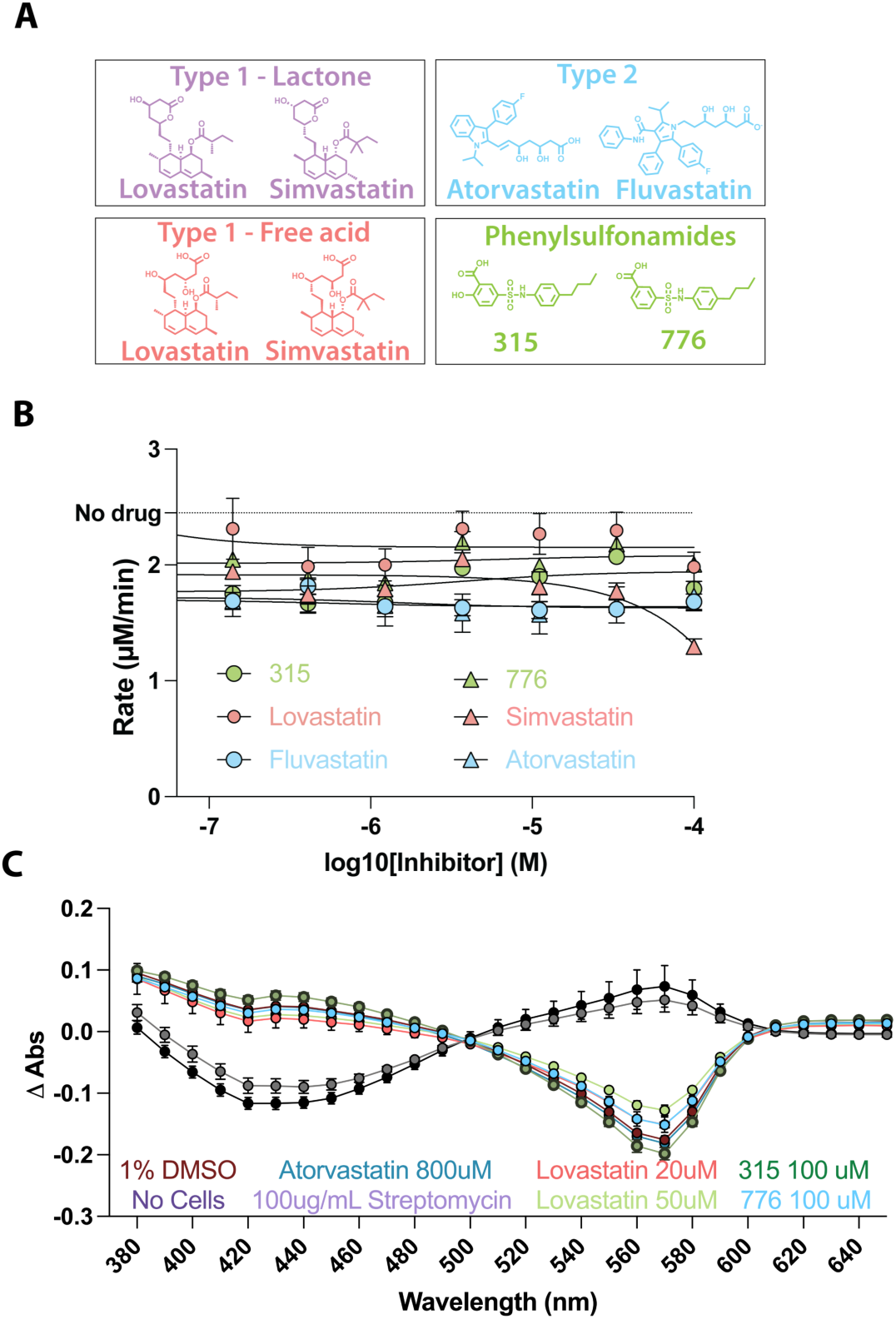
Assessment of inhibitors targeting *B. burgdorferi* HMGR. **A**) Chemical structures of Type 1 and 2 statins and phenyl sulfonamides that inhibit the *E. faecalis* HMGR. **B**) Dose response for statins and phenyl sulfonamides 315 and 776 against purified *B. burgdorferi* HMGR. Assays were performed in triplicate using an absorbance-based assay and fit to log(inhibitor) versus response. **C**) Absorbance spectra of *B. burgdorferi* B31 K2 cultures (∼1×10^7^ cells/mL) grown in the presence of DMSO (diluent), streptomycin (known growth inhibitor), lovastatin at indicated concentrations (red and green), or atorvastatin (blue) using the pH indicator phenol red. Absorbance was measured every 10 nm from 350 to 650 nm. Absorbance values from day 0 were subtracted from day 8 to get Δ Abs which was plotted against the wavelength. Shown are the means and standard deviations for three technical replicates.

**Fig. S8.**
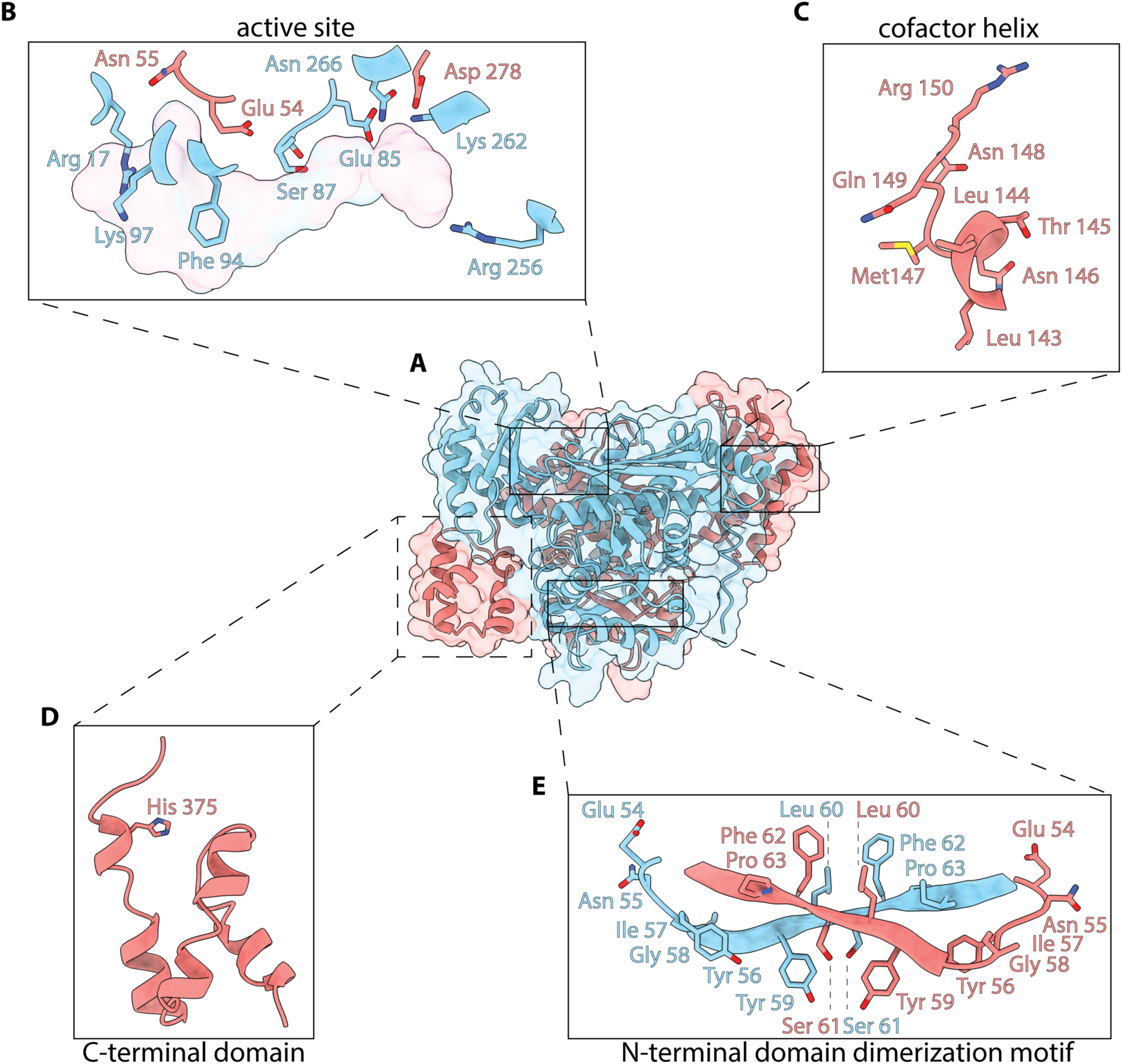
Crystallographic characterization of *B. burgdorferi* HMGR: **A**) The homodimeric structure is depicted in both cartoon and a transparent surface format. One monomer is colored in blue while the other is shown in salmon. Several key functional domains are highlighted. **B**) Active site residues involved in binding substrate. **C**) Residues on the cofactor helix that engage with NAD(P)H in other bacterial HMGRs. **D**) The C-terminal domain with the catalytic His 375 shown in stick format. **E**) Residues that comprise the dimerization motif.

**Fig. S9.**
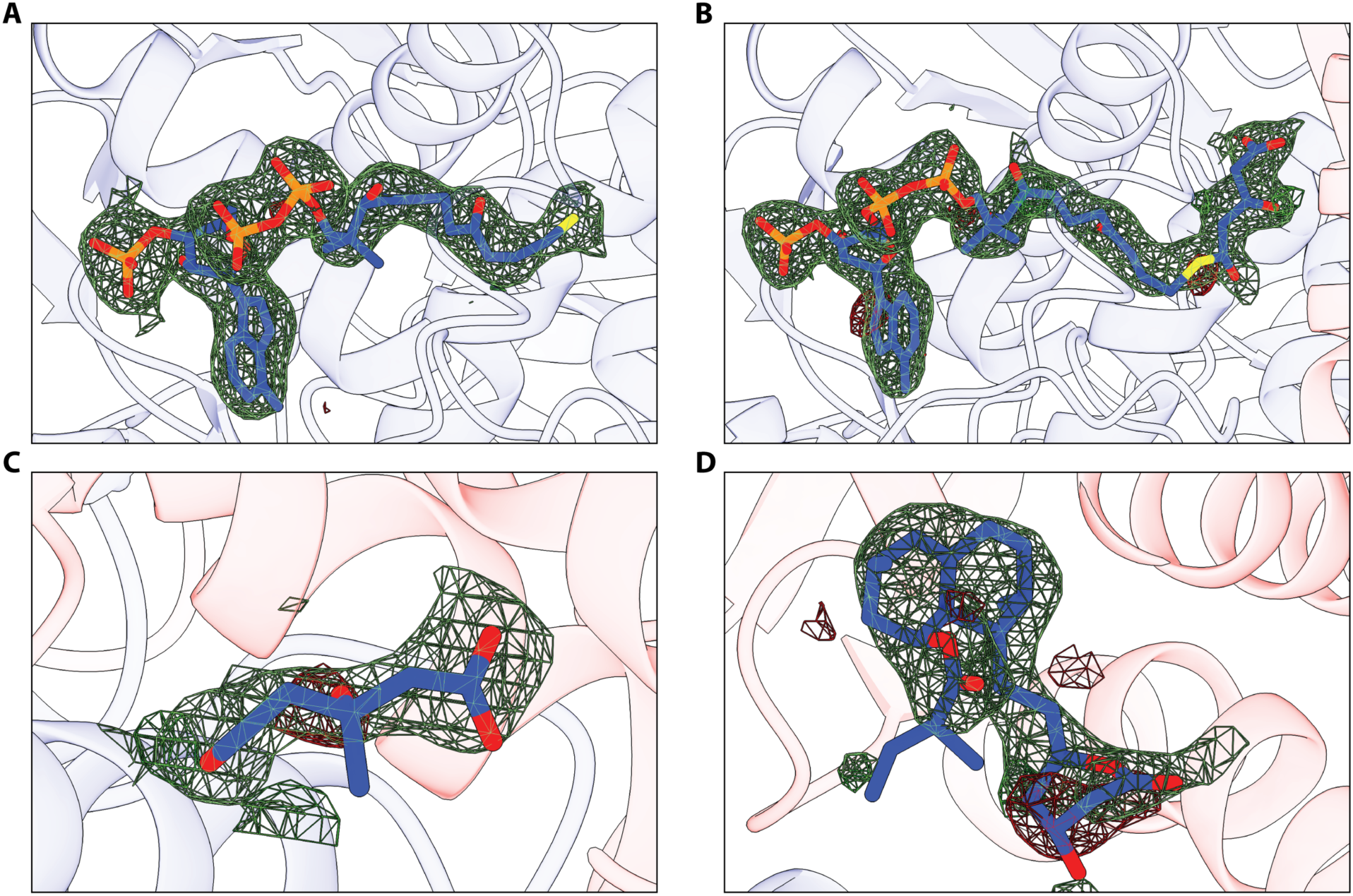
Omit densities for *B. burgdorferi* HMGR ligands. mFo-DFc omit maps contoured at 3σ for **A**) CoA, **B**) HMG-CoA, **C**) mevalonate, and **D**) lovastatin bound to *B. burgdorferi* HMGR.

## Supporting tables

**Table S1.**
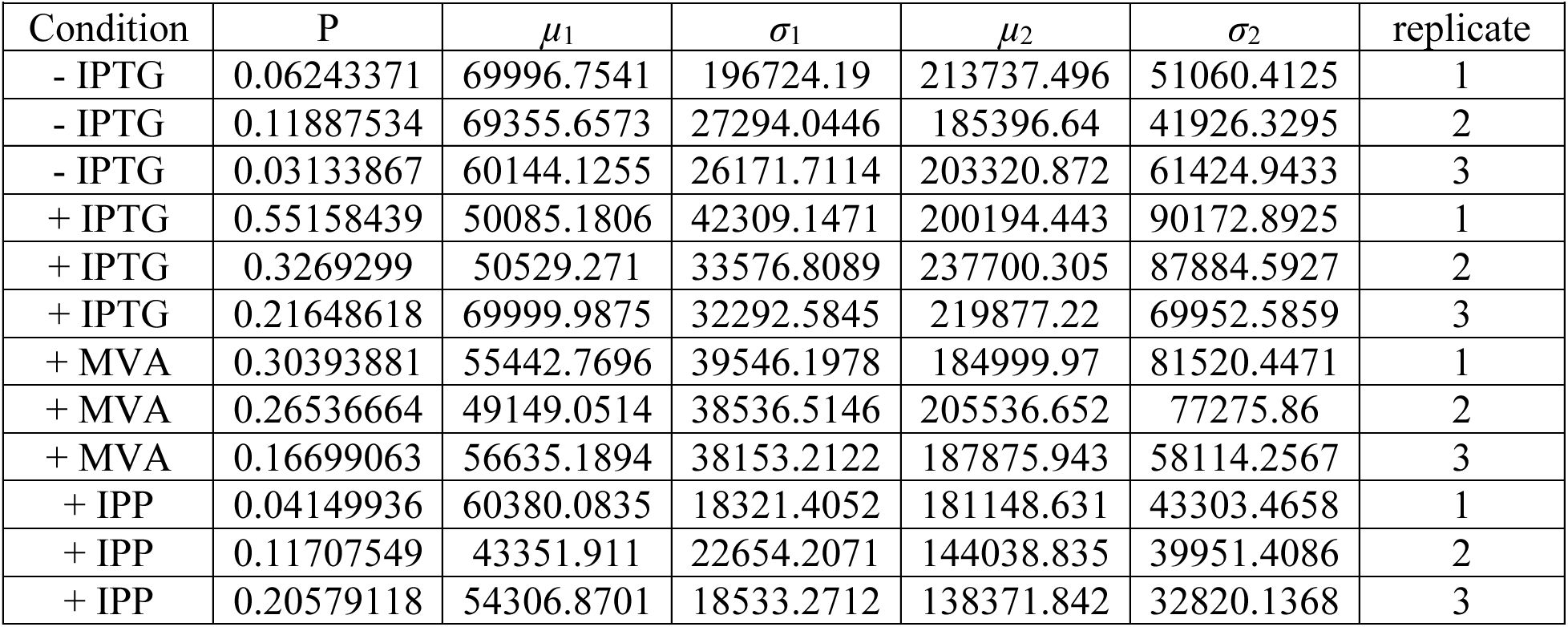
Fitting results for single cell analysis for the HADA labeling experiments.

|  |  |  |  |  |  |  |
| --- | --- | --- | --- | --- | --- | --- |
| Condition | P | $\mu_1$ | $\sigma_1$ | $\mu_2$ | $\sigma_2$ | replicate |
| - IPTG | 0.06243371 | 69996.7541 | 196724.19 | 213737.496 | 51060.4125 | 1 |
| - IPTG | 0.11887534 | 69355.6573 | 27294.0446 | 185396.64 | 41926.3295 | 2 |
| - IPTG | 0.03133867 | 60144.1255 | 26171.7114 | 203320.872 | 61424.9433 | 3 |
| + IPTG | 0.55158439 | 50085.1806 | 42309.1471 | 200194.443 | 90172.8925 | 1 |
| + IPTG | 0.3269299 | 50529.271 | 33576.8089 | 237700.305 | 87884.5927 | 2 |
| + IPTG | 0.21648618 | 69999.9875 | 32292.5845 | 219877.22 | 69952.5859 | 3 |
| + MVA | 0.30393881 | 55442.7696 | 39546.1978 | 184999.97 | 81520.4471 | 1 |
| + MVA | 0.26536664 | 49149.0514 | 38536.5146 | 205536.652 | 77275.86 | 2 |
| + MVA | 0.16699063 | 56635.1894 | 38153.2122 | 187875.943 | 58114.2567 | 3 |
| + IPP | 0.04149936 | 60380.0835 | 18321.4052 | 181148.631 | 43303.4658 | 1 |
| + IPP | 0.11707549 | 43351.911 | 22654.2071 | 144038.835 | 39951.4086 | 2 |
| + IPP | 0.20579118 | 54306.8701 | 18533.2712 | 138371.842 | 32820.1368 | 3 |

**Table S2:**
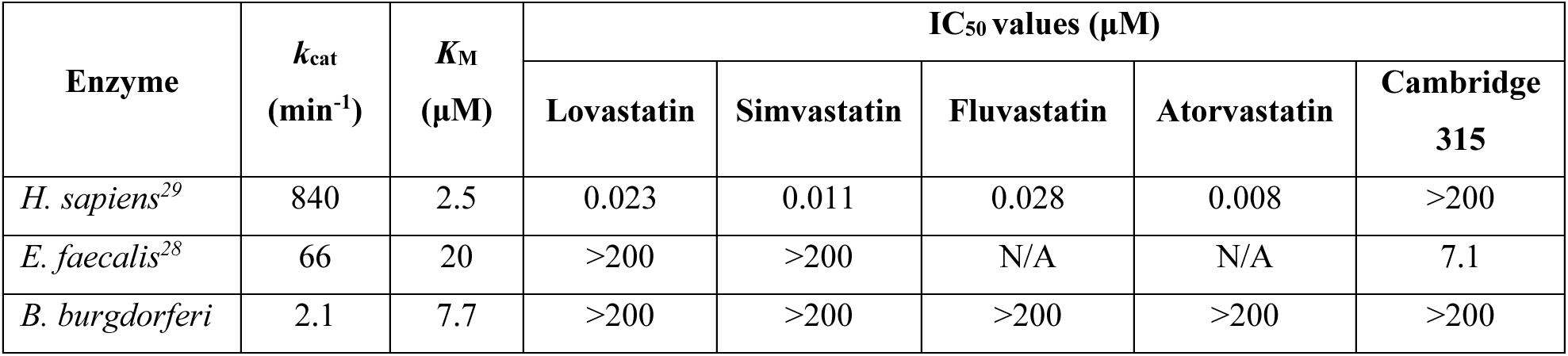
Steady-state kinetics and inhibition of bacterial and human HMGRs.

**Table S3:**
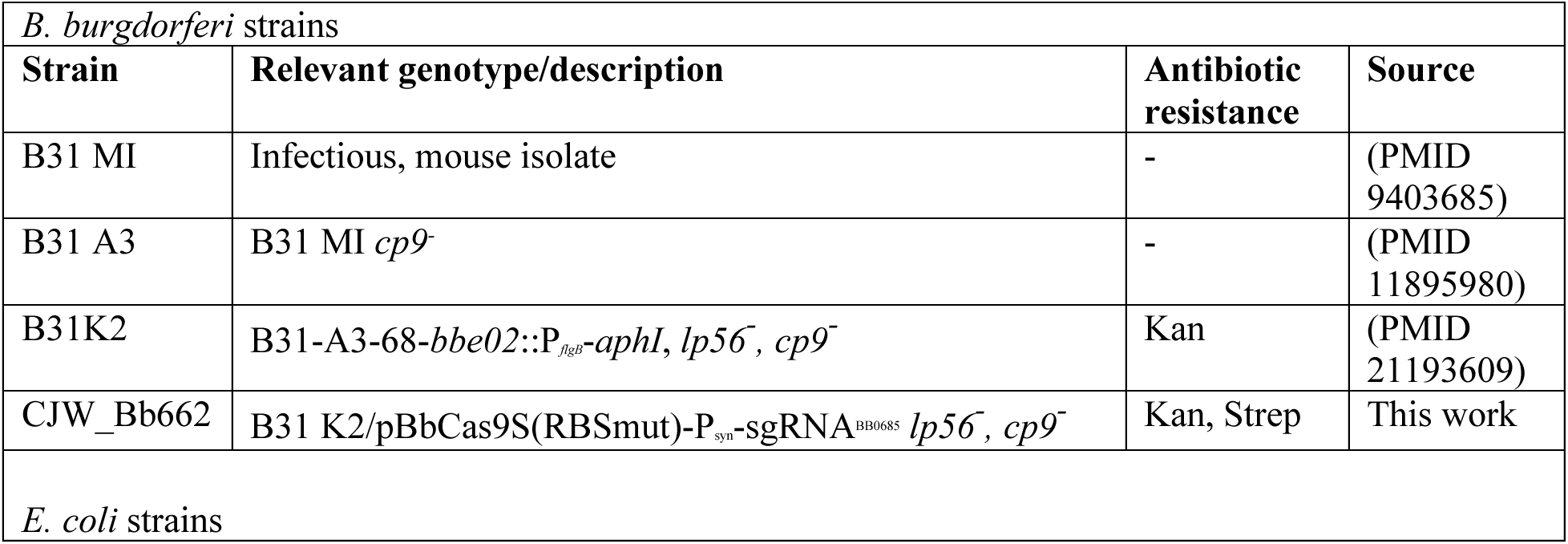
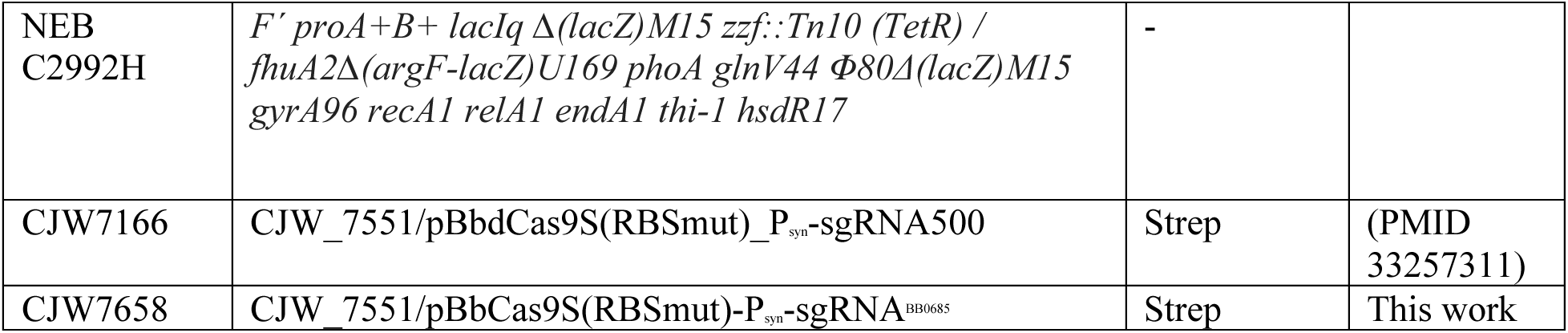
Bacterial strains used in this study.

**Table S4.** Bacterial plasmids used in this study.

| Table S4. Bacterial plasmids used in this study |  |  |  |
| --- | --- | --- | --- |
| Plasmid | Description | Antibiotic resistance | Source |
| pBbdCas9S(RBSmut)_P<sub>syn</sub>-sgRNA500 | Parent for the “all-in-one” CRISPRi shuttle vector. It has a mutation in the RBS to help reduce leakiness of the dCas9 expression. | Strep | (PMID 33257311) |
| pBbCas9S(RBSmut)-P<sub>syn</sub>-sgRNA<sup>BB0685</sup> | “All-in-one” shuttle vector that includes an sgRNA targeted against *bb0685*. | Strep | This work |

## Protein sequences

*Borreliella burgdorferi* (strain ATCC 35210 / DSM 4680 / CIP 102532 / B31) Uniprot: O51628

MNLESLSSFMELSKNFRHKSVLEKRQEIKSFLELSYKDFFYNNANEDFLFNMIENYIGYL SFPIGIVKNLKINGKYYSLPIATEESSVVAALNFAAKILENADLRYSLGEVLGISQIYIKSE KDLSKIFVDLGDKIKTWIEPLLTNMNQRGGGFRRLSTRHIKELGIQKLNIYVDTCDAMGA NLLNSIAERVAEFIFLEFGYECVLKVLSNDISEFTAKARFVLDFKHLLPGKEDSWNLAKKI ELISSIGFYEEERAVTNNKGIMNGITGVCLATFNDTRALEASVHKFASKSGKYFPLSKFYT TDNALVGEIEIPLQVG TKGGVISFNEASILSFKIMNVNSKSEFIGILSCVGLASNFAALRALAFNGIQKGHMRLHVN KILHLLKTKYNISDFEKDKLLLEMERMNIYSFDFAFKILKKIRLENENKV

## Lactobacillus jensenii

Uniprot: A0A5N1IG17

MKFYQLPISERRKMLLQNGIKLNHVDDDLLSELDLLSENVIGKLTLPLSVLQTAIVNGQS FQVPMATEESSVVAAANHGLNIFNQNGGVSAKSERTGIWGQLVFEVAEFSLAEFEAKKP DYLKLVNEEFASLVKHGGGVRQIIAEVKTDLLFLRVLVDPAESMGANRTNTILEFLGQKI SQDFTIEKLYAILSNYPSQYTCAKVSLAFASLTKTKDEKIGEKIAQKIVLLSKIGQEDPYR AVTNNKGIMNGVDAILLATGNDFRAVEAACHQAASLSGSYQSLSNWRIEDNKLVGEIK LPLAIGVVGGSIKSRSDVQVAYRILRQVTASELAEIIAAVGLANNLAALLAISTVGIQKGH MSLQIRNVLKNLTATDEEKNSVKELMQKQKRYSETDAKKFLQEIREENN

## Flavobacterium psychrophilum

Uniprot: A0A076P6Q4

MPKLTTGFSKLSKEEKINWIASTHFSNAEEATQTIKKYWNSDLELQKLHDEFIENTITNFYLPLGVAPNFLINGKNHTIPFAIEESSVVAAASKSAKYWGTRGGFKTTVLNSEKIGQVHFIFKGDSRNLIVFFNQIKSKLFAHTESITTNMQKRGGGILDIELRDKTSDLENYYQLHATFETKDSMGANFINSCLEQFAKTLKEEALQSEILSETEKNIEVVMSILSNYVPNCIVRAEVSCPVSDLSEKNIENPQEFAQKFIRAVKIAEIEPFRAVTHNKGIMNGIDAVVLATGNDFRAVEAGIHAYAARNGQYSSLSHAKIENDIFTFWLEIPLALGTVGGLTSLHPLVKMSLEMLEKPSAQELMQIVAVAGLAQNFAALRSLTTTGIQQGHMKMHLNNIINQFEATQNERVLIKNHFTENTVSHSAVVAFIESLRK

*Legionella pneumophila subsp. pneumophila* (strain Philadelphia 1 / ATCC 33152 / DSM 7513) Uniprot: Q5ZTV6

MSIAANASELFRGFSKLSREERFQRLCALGALTHEDITFLKQGGIKDLNLADKLIENVIGYFQLPLGVATNFNIDGRDYVIPLAVEETSIIAALSKSAKWIRQHGEINTWVHGECILGQIQLAKVKDFQRFSDLFNKNRQYFIEIANKDVAANMVKRGGGVTDLQVRHLKREDGLDMAVIHLTMNSCDAMGANIINQVLEYLKQPIEQITGEEVTMCILSNLNDQKLTTAQVIIRNIDPILGQKLQEASLFAEIDPYRAATHNKGVMNGIDPVLIATGNDWRAVEAGIHAYAARSGQYKAITRWRYQNEILTGELTAPIIVGTVGGVTSLHPTAKMCLRMMDITSANQLSQVIAAVGLVQNLGALKALCTDGIIQGHMKLHIDNLLLVAGANENEMPVLKEKLQEWLNLNKRVSLNN AYDLLAEIRQAPVAV

